# A subset of SMN complex members have a specific role in tissue regeneration via ERBB pathway-mediated proliferation

**DOI:** 10.1101/678417

**Authors:** Wuhong Pei, Lisha Xu, Zelin Chen, Claire C Slevin, Kade P Pettie, Stephen Wincovitch, NISC Comparative Sequencing Program, Shawn M Burgess

## Abstract

Spinal Muscular Atrophy (SMA) is the most common genetic disease in childhood. SMA is generally caused by mutations in *SMN1*. The Survival of Motor Neurons (SMN) complex consists of SMN1, Gemins (2–8) and Strap/Unrip. We previously demonstrated *smn1* and *gemin5* inhibited tissue regeneration in zebrafish. Here we investigated each individual SMN complex member and identified *gemin3* as another regeneration-essential gene. These three genes are likely pan-regenerative since they affect the regeneration of hair cells, liver and caudal fin. RNA-Seq and miRNA-Seq analyses reveal that *smn1, gemin3*, and *gemin5* are linked to a common set of genetic pathways, including the tp53 and ErbB pathways. Additional studies indicated all three genes facilitate regeneration by inhibiting the ErbB pathway, thereby allowing cell proliferation in the injured neuromasts. This study provides a new understanding of the SMN complex and a potential etiology for SMA and potentially other rare unidentified genetic diseases with similar symptoms.

## Introduction

Spinal Muscular Atrophy (SMA) is the leading hereditary cause of infant mortality (1, 2). The majority of SMA cases are caused by mutations in the *survival of motor neuron 1* (*SMN1*) gene. *SMN1* is ubiquitously expressed and a reduction of SMN1 protein leads to motor neuron death in patients afflicted with SMA. Although the incidence of SMA is approximately 1:6000 in live births, the carrier frequency for a heterozygous *SMN1* mutation can approach 1:40 in adults. Many important questions remain regarding the pathology of the disease, including why the ubiquitously expressed SMN1 protein primarily impacts motor neurons, which other organs are potentially affected by SMN1 deficiencies, and whether SMA is a developmental or degenerative disease (or both).

The SMN1 protein is part of the SMN complex, responsible for the assembly of small nuclear ribonucleoproteins (snRNP) needed for pre-mRNA splicing (3, 4). The SMN complex consists of nine proteins, however the majority of research on the complex has focused on the characterization of SMN1 and its role in SMA. In addition to its role in the SMN complex, SMN1 plays a role in many other biological processes, including axon growth, mRNA transport, and translational control (5, 6). Although there is some evidence showing that other SMN complex members, such as GEMIN3 and GEMIN5, also have functions independent of the SMN complex (7, 8), it remains largely unknown how the other SMN complex members relate to SMA, and whether other members have functions beyond the SMN complex. A reduction of the SMN1 protein in SMA results in the reduction of other SMN complex members (9), suggesting that there is a functional inter-dependence among the nine genes.

In a previous study (10), we showed that mutations in *smn1* and *gemin5* negatively impacted the ability of zebrafish to regenerate different tissues after injury. Regenerative medicine is a rapidly expanding field of science that focuses on replacing or regenerating organs damaged by injury, aging, or degenerative conditions. An active area of research within regenerative medicine is the restoration of hearing by replacing the lost mechanosensory receptors of the inner ear known as hair cells. Age-related hair cell death impairs the hearing of hundreds of millions of the elderly and hearing loss as a side-effect of therapeutic medications remains a major concern (11). In general, mammals have very limited regenerative capability, however many non-mammal animal models including the zebrafish have been used extensively because they possess a much broader capacity for wound healing, including the capacity to regenerate hearing after damage. Zebrafish are particularly well suited for studying the regeneration of hair cells because of a second organ that fish and amphibians possess on their skin known as the lateral line, which consists of clusters of hair cells in structures known as “neuromasts” (12). Similar to the case in the mammalian inner ear, hair cells in the neuromasts are surrounded by support cells, which in fish and amphibians can replace the lost hair cells through either mitotic division or trans-differentiation. Support cells in the zebrafish neuromast are further surrounded by mantle cells, which resemble quiescent stem or progenitor cells (13). Hair cell regeneration studies in zebrafish have uncovered numerous genetic factors and molecular pathways that are associated with the regeneration of hair cells (14); conversely, random mutagenesis studies revealed that mutations in only a small number of genes actually significantly alter hair cell regeneration specifically (10, 13, 15, 16). There is consistently a gap between the number of genes transcriptionally associated with regeneration and the number of genes essential for regeneration in other tissues as well (17, 18). Therefore, seeking novel genes essential for tissue regeneration is pivotal in understanding the core molecular mechanisms of wound healing, and for ultimately advancing regenerative medicine.

In this study, we systemically mutated all nine genes encoding SMN complex proteins. Using hair cell regeneration in the zebrafish lateral line as an assay, we identified three SMN complex members as essential factors that regulate regeneration through ErbB pathway-mediated cell proliferation. Additional studies revealed that these regenerative members were also essential for the regeneration of other tissues and all shared common transcriptional pathways altered in the mutant larvae. Our findings demonstrated a subset of the SMN complex proteins had separate functional roles involved in tissue regeneration.

## Results

### Divergent roles for SMN complex members in embryo development and hair cell regeneration

Hearing loss is one of the common sensory disorders negatively affecting the quality of life for hundreds of millions of people worldwide (11). In a search for novel genes involved in hearing regeneration, we previously performed a large-scale mutagenesis screen in zebrafish and identified *smn1* and *gemin5* as essential genes for hair cell regeneration (10). Both Smn1 and Gemin5 belong to the SMN complex, a multiprotein complex functioning in the biosynthesis of small nuclear ribonucleoproteins (snRNP). To investigate whether the regenerative abilities of Smn1 and Gemin5 are linked to the SMN complex activity, we independently mutated all nine genes in the SMN complex (Suppl. Table 1) and examined their involvement in hair cell regeneration. We found that in addition to *smn1* and *gemin5*, mutations in *gemin3* altered hair cell regeneration but had no effect on initial hair cell development (Fig. 1A. Suppl. Fig. 1A. We also found that mutations in the other six SMN complex genes, *gemin2, gemin4, gemin6, gemin7, gemin8* and *strap/unrip*, had no impact on hair cell regeneration (Fig. 1D-I). None of the nine mutants showed an overt morphological phenotype in early larvae (data not shown), but all mutants except *gemin8* and *strap* failed to survive to adulthood (Suppl. Table 2). Altogether, these data classified the functions of the nine members of the SMN complex into three categories: three (Smn1, Gemin3 and Gemin5) are essential for hair cell regeneration and adult survival; four (Gemin2, Gemin4, Gemin6 and Gemin7) are essential for adult survival but not for hair cell regeneration; two (Gemin8 and Strap) are required for neither.

**Fig. 1.**
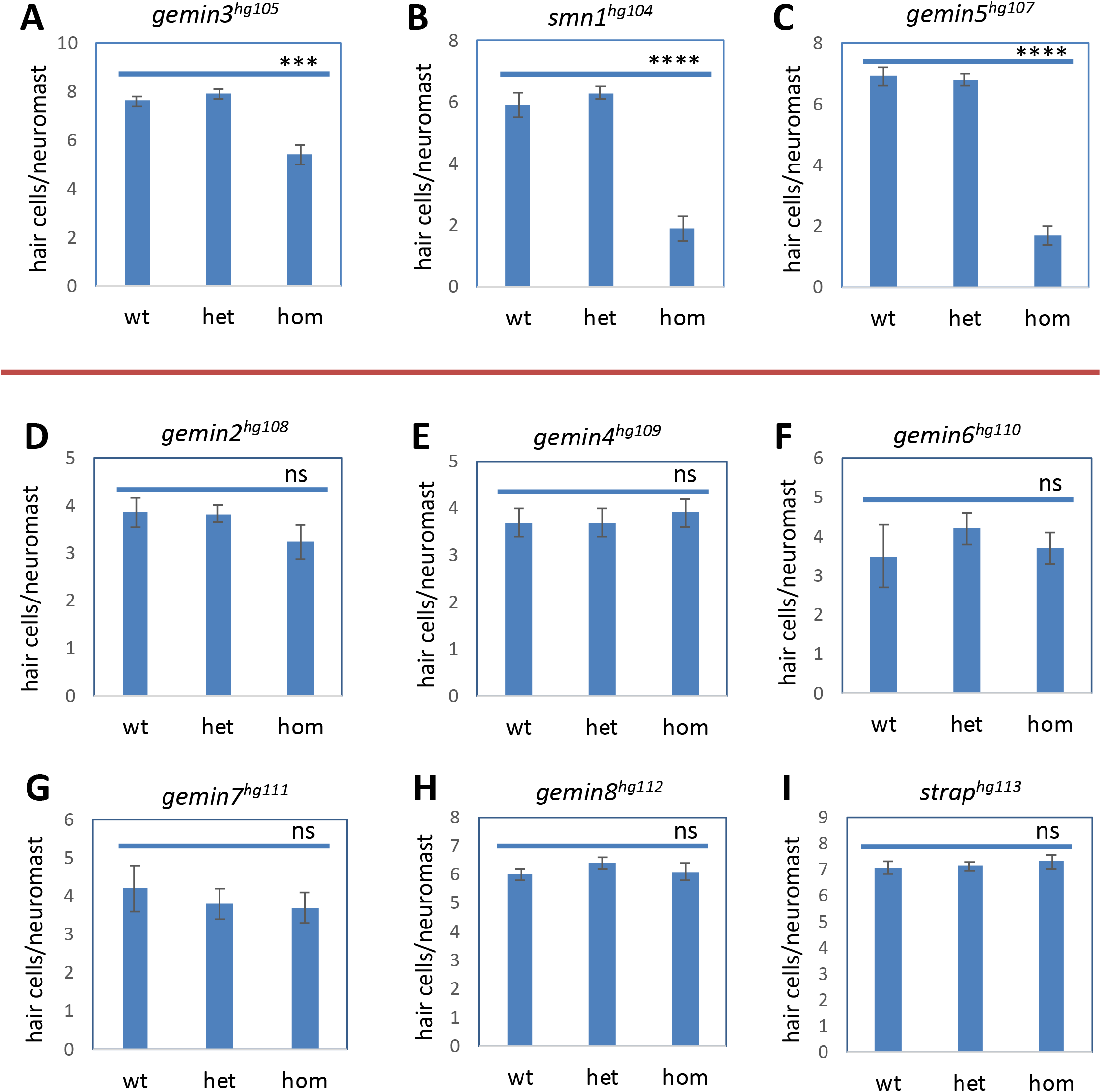
Hair cell regeneration analysis for mutations in each gene of the SMN complex. Hair cell regeneration is impaired by homozygous mutations of *gemin3^hg105^* (A), *smn1^hg104^* (B) and *gemin5^hg107^* (C), but not in *gemin2^hg108^* (D), *gemin4^hg109^* (E), *gemin6^hg110^* (F), *gemin7^hg111^* (G), *gemin8^hg112^* (H) or *strap^hg113^* (I). Red line separates the mutations impacting regeneration from those that have no effect on regeneration. wt, wild-type. het, heterozygotes. hom, homozygotes. Error bars in the graphs represent mean ± s.e.m. The difference between wild-type and homozygote is labeled. ns, P > 0.05; ***P < 0.001; ****P < 0.0001.

We generated an additional mutation allele for the three genes involved in regeneration, *smn1, gemin3*, and *gemin5* to verify their role in hair cell regeneration. The second mutation alleles all recapitulated the deficits in hair cell regeneration (Fig. 1B-C, Suppl. Fig. 1B). To examine whether the regeneration was specific to the ablation of hair cells using CuSO_4_, we performed the ablation using neomycin and observed similar regeneration deficits (Suppl. Fig. 1C-E).

In support of their divergent phenotypes in hair cell regeneration and adult survival, whole-mount in situ hybridization analysis showed that the SMN complex genes possessed some common but also some different expression patterns at 3 days post fertilization (dpf) (Suppl. Fig. 2A-H). The brain expression of these genes in particular revealed both shared and specific expression patterns: five of these genes were restricted to a stripe at approximately the mid-hindbrain boundary. *smn1* was more enriched in the eye regions and *gemin5* was more condensed at the midline of the brain; *gemin3* expression was relatively weaker than the others; *gemin7* and *strap* showed a ubiquitous expression which was different from the other gemins. At 1 dpf, a stronger similarity was detected in the brain expression between *smn1* and *gemin5* (Suppl. Fig. 2I). Both genes were enriched in the eyes, brain and midline area.

Our whole-mount in situ hybridization analysis showed that none of the SMN complex genes were particularly enriched in the lateral line neuromasts (data not shown). However, single cell RNA-sequencing analysis conducted by the Piotrowski group demonstrated that all these SMN complex genes are expressed at detectable levels in lateral line neuromasts, and different genes in the complex are expressed in different neuromast cell types at different levels (Suppl. Fig. 2J) (13). The non-identical and complex patterns of gene expression for the different SMN subunits (as well as the different phenotypes) suggest that the various roles for each protein may not be simply as co-expressed subunits, but that the composition of the SMN complex and potentially alternate functions of the subunits may vary based on expression levels and cellular context.

Maternal mRNA and protein deposition allows zebrafish embryos to grow rapidly during the first few hours after birth and some maternal proteins can persist for days after fertilization. Although regeneration was analyzed at 7 dpf, we still examined whether the hair cell regeneration phenotype could be associated with the initial maternal deposition or a difference in the stability of mutant mRNAs. We analyzed the expression level of two regeneration genes (*smn1* and *gemin5*) and three non-regeneration genes (*gemin4, gemin6* and *strap*) at different stages of embryonic development by semi-quantitative PCR and found no clear difference between these two groups of genes (Suppl. Fig. 2K), suggesting mRNA destabilization does not explain hair cell regeneration phenotypes or eventual larval death.

Genetic interactions have been observed among SMN complex genes (19, 20). To study whether there is a synergy among the three genes involved in regeneration, we generated an *smn1* and *gemin5* double mutant and studied the effect of simultaneous depletion of two genes on morphology and hair cell regeneration. We found the *smn1* and *gemin5* double mutant had a normal embryonic morphology and normal hair cell development (data not shown), as observed in both the *smn1* and *gemin5* single mutants. The double mutant showed the expected deficiency in hair cell regeneration; however, the level of deficiency was comparable to that of the *gemin5* mutant (Suppl. Fig. 3A-C). Taken together, these data suggest there is no functional synergy among these regeneration genes, and *smn1* and *gemin5* appear to both be necessary and fall in the same regenerative pathway as the phenotypes in double mutants were neither synergistic nor additive.

### The overall neuromast size is smaller in mutants with regenerative phenotypes

We examined the neuromast cell patterning in the mutants and control siblings at 2 days post hair cell ablation to see if we could detect a difference in neuromast size in mutants using both transgenic labeling and immunohistochemical staining approaches. Double transgenic labeling of support cells by Tg(*tnks1bp1*:EGFP) and hair cells by Tg(*atoh1a*:dTomato) in *gemin5* mutants revealed that support cells in the mutant occupied a reduced area likely because of fewer cells and hair cells were fewer when compared to that of the control siblings (Fig. 2A). Whole neuromast labeling using Tg(*cldnb*:EGFP) showed that the size of the neuromast in the mutant was smaller than that of the control siblings (Fig. 2B). Alkaline phosphatase staining revealed that the structure of lateral line neuromasts were much more reduced in the mutant (Fig. 2C). Co-staining with anti-hair cell antibody and nuclear dye DAPI revealed a reduction in the number of hair cells and neuromast cells (Fig. 2D-E).

**Fig. 2.**
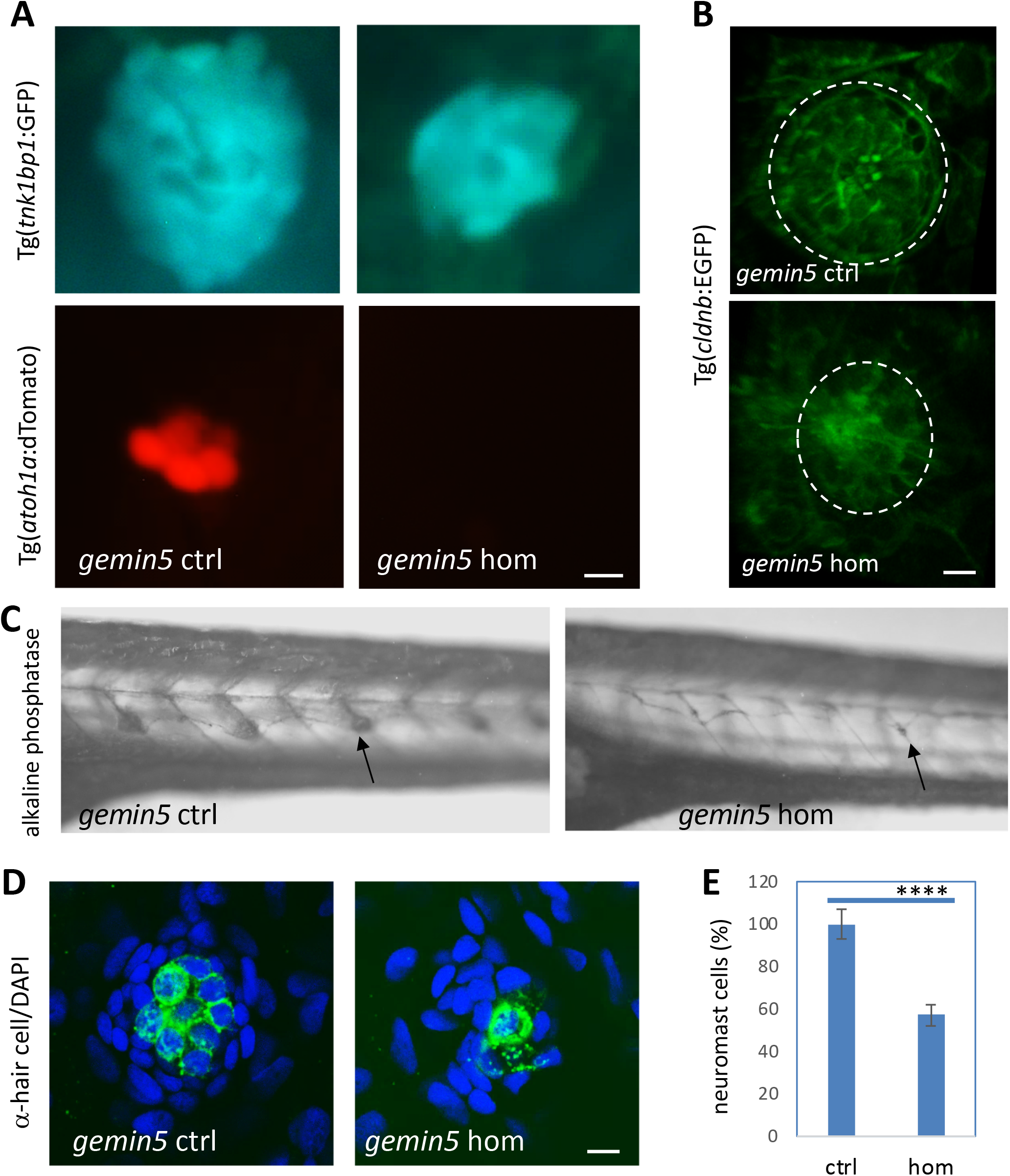
*gemin5^hg81^* mutants possess smaller neuromasts at 2 days post hair cell ablation. (A) Live cell imaging of support cells by Tg(*tnks1bp1*:GFP) and hair cells by Tg(*atoh1a*:dTomato) in lateral line neuromasts of the *gemin5^hg81^* control and mutant embryos at 2 days post hair cell ablation. Scale bar, 10 μm. (B) Live cell labeling of neuromast cells by Tg(*cldnb*:EGFP) in the lateral line of the *gemin5^hg81^* control and mutant embryos at 2 days post hair cell ablation. Dotted white circle demarcates the periphery of the neuromast. Scale bar, 10 μm. (C) Alkaline phosphatase staining of lateral line neuromasts in the *gemin5^hg81^* control and mutant embryos at 2 days post hair cell ablation. Arrows point to the neuromasts. (D) Confocal images of lateral line neuromasts in the *gemin5^hg81^* control and mutant embryos at 2 days post hair cell ablation, stained with anti-hair cell antibodies (green color) and DAPI (blue color). Representative images are shown. Scale bar, 10 μm. (E) Quantification of neuromast cells. Error bars in the graphs represent mean ± s.e.m. There is a significant reduction in the number of neuromast cells (**** P < 0.0001). The numbers are presented as percentage because they were obtained from quantification of still confocal images.

We also used transgenes and immunohistochemical staining to examine the neuromast cells at 2 days post hair cell ablation in *smn1* and *gemin3* mutants. Consistent with the results of the *gemin5* mutation, mutations in *smn1* and *gemin3* also caused a reduced area of support cells, impaired regeneration of hair cells and smaller neuromasts (Suppl. Fig. 4). All the data suggest the proliferative capacity in the neuromasts is reduced preventing organ growth.

### Regenerative deficient mutants show reduced proliferation after injury

To directly test proliferative capacity of the support cells in the mutant neuromasts, we used an EdU incorporation assay to label the proliferation of neuromast cells after hair cell ablation. Compared to the control siblings, all three mutants possessed a reduced number of EdU positive cells (Fig. 3), suggesting that after hair cell ablation, the mutants lack the capacity to effectively proliferate either their support cells or mantle cells.

**Fig. 3.**
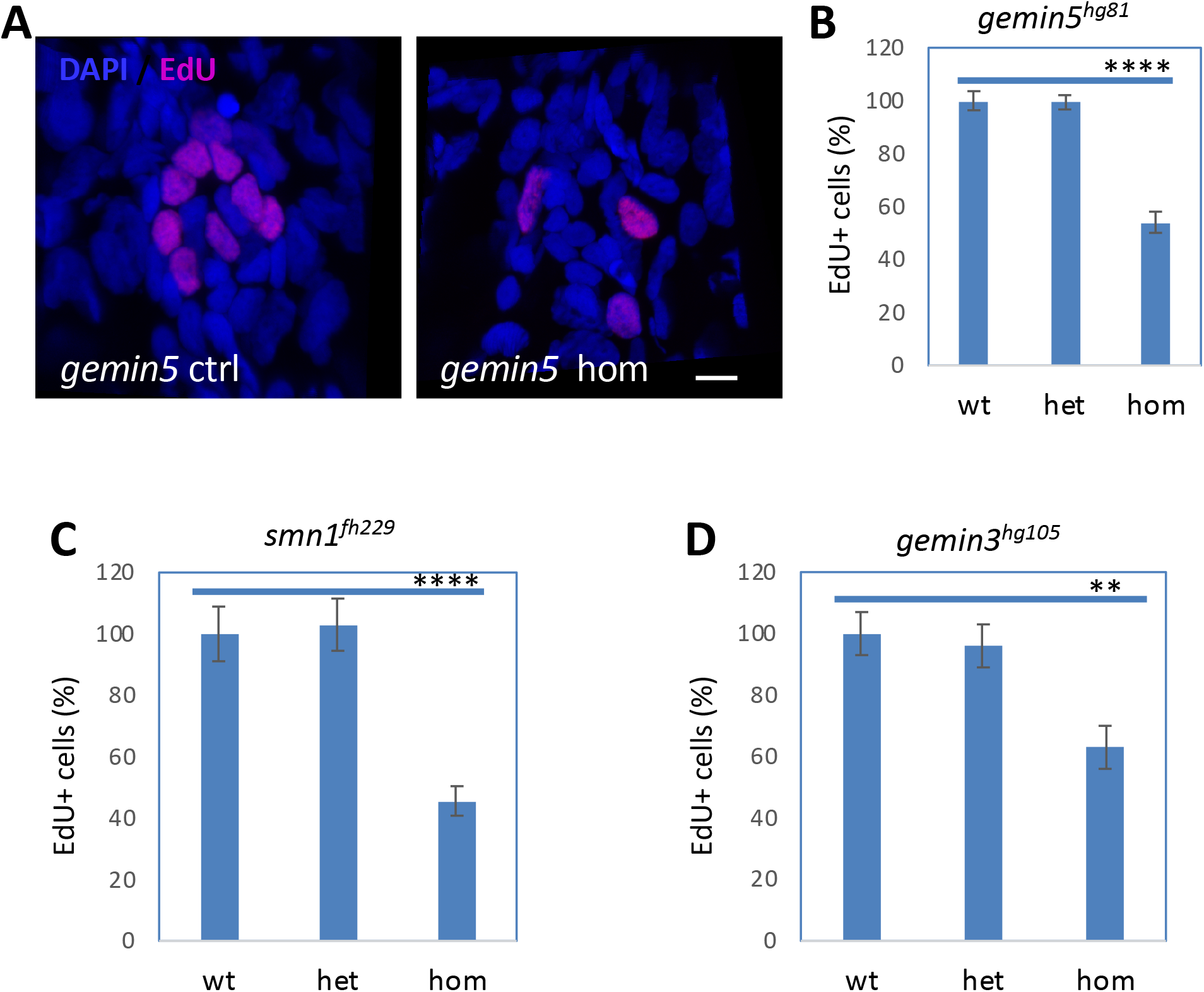
Decreased neuromast cell proliferation after hair cell ablation in *gemin3, gemin5*, and *smn1* mutants. (A) Confocal images of lateral line neuromasts in the control and *gemin5^hg81^* mutant at 1-day post hair cell ablation. Neuromasts were stained with DAPI (blue) and proliferating cells were labeled by EdU (pink). The embryos used for the analysis were obtained from a pairwise incross of heterozygotic parents. Hair cells were ablated at 5 dpf. EdU treatment was conducted at 1-day post ablation. Scale bar, 10 μm. (B) Quantification of EdU positive cells in the embryos carrying wild-type, heterozygous or homozygous *gemin5^hg81^* mutations. (C) Quantification of EdU positive cells in the embryos carrying wild-type, heterozygous, or homozygous *smn1^fh229^* mutations. (D) Quantification of EdU positive cells in the embryos carrying wild type, heterozygous or homozygous *gemin3^hg105^* mutations. The graphs show mean ± s.e.m. Homozygous mutants for all three regeneration genes have a significantly reduced number of proliferating cells. **P < 0.01; ****P < 0.0001.

### Regeneration deficient mutants are less sensitive to the ErbB pathway inhibitor AG1478

We conducted numerous tests to find pathways affected by *gemin5* mutations. Most conditions were negative (Suppl. Table 3), with only AG1478 (inhibiting ErbB signaling) showing a specific phenotype. Treatment with 2 μM AG1478 caused a dramatic increase in lateral line neuromasts of control siblings, but only a mild increase in the *gemin5* mutant larvae (Fig. 4A-B), indicating that the *gemin5* mutant is resistant to ErbB pathway inhibition. To determine whether the reduced sensitivity was common to all three regeneration gene mutations, we also treated *smn1* and *gemin3* mutants with AG1478. Consistent with the findings from the *gemin5* mutant, the *smn1* mutant and the *gemin3* mutants also showed a reduced responsiveness to AG1478 (Fig. 4C and D). To test if the ErbB pathway responded normally in other mutants in the complex, we treated *gemin6* mutants with AG1478. In contrast to the mutants that disrupted regeneration, the *gemin6* mutant responded to AG1478 comparable to that of their control siblings (Fig. 4E). Altogether, these results point out that the inability to respond to AG1478 inhibition specifically in the mutants that inhibited regeneration, suggesting a mechanistic link.

**Fig. 4.**
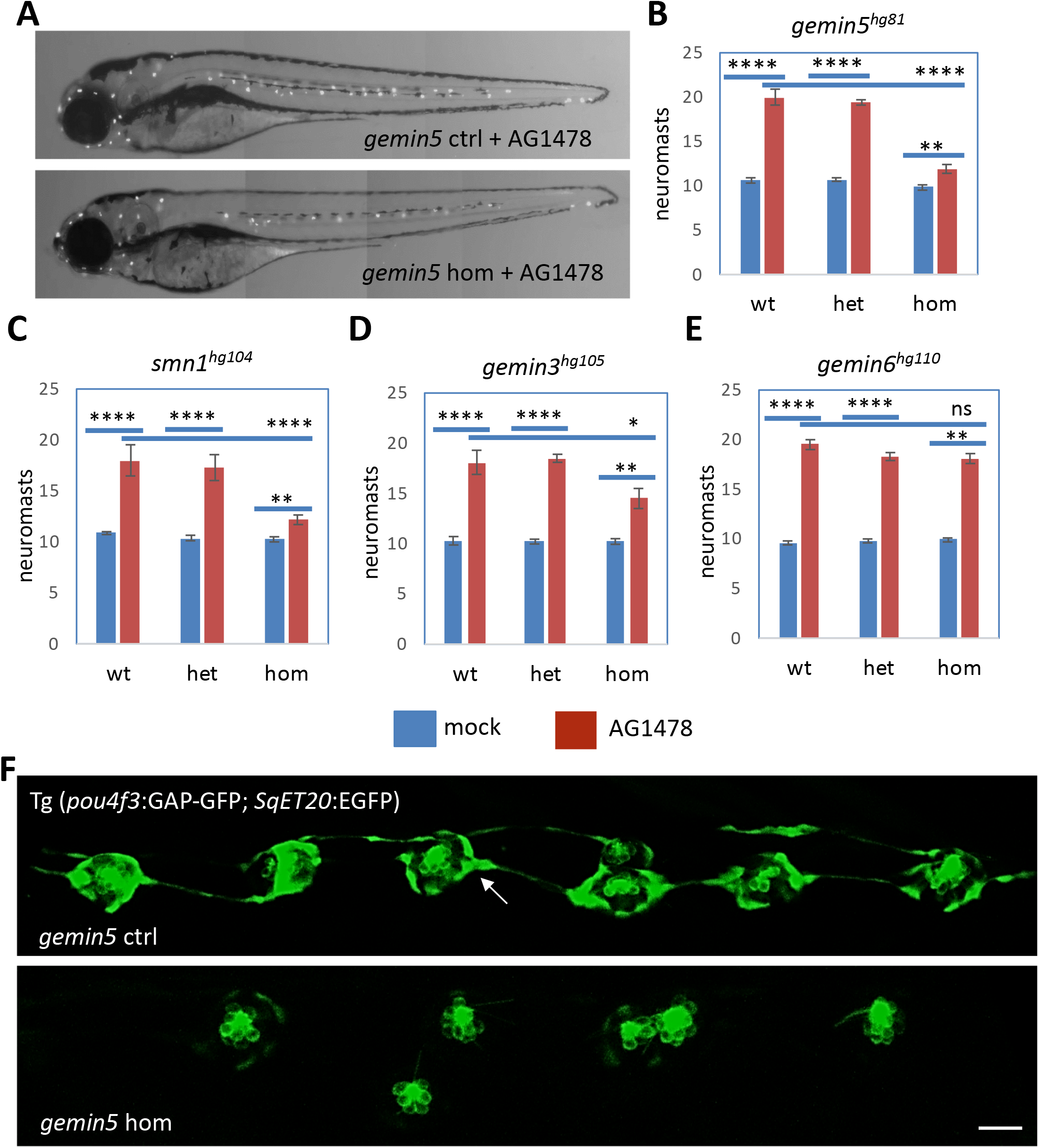
Reduced responsiveness of *gemin3, gemin5*, and *smn1* mutants to ErbB pathway inhibitor AG1478. (A) Neuromasts in AG1478-treated control and *gemin5^hg81^* mutant embryos at 5 dpf. Neuromasts are shown as white dots. (B) Quantification of the lateral line neuromasts in mock and AG1478 treated embryos carrying the *gemin5^hg81^* mutation at 5 dpf. (C) Quantification of the lateral line neuromasts in mock and AG1478 treated embryos carrying the *smn1^hg104^* mutation at 5 dpf. (D) Quantification of the lateral line neuromasts in mock and AG1478 treated embryos carrying the *gemin3^hg105^* mutation at 5 dpf. (E) Quantification of the lateral line neuromasts in mock and AG1478 treated embryos carrying the *gemin6^hg110^* mutation at 5 dpf. Error bars in the graphs show the mean ± s.e.m. ns, P > 0.05; *P < 0.05; **P < 0.01; ****P < 0.0001. (F) Fluorescent images of lateral line neuromasts labeled by transgenes Tg(*pou4f3*:GAP-GFP) and Tg(*SqET20*:EGFP) in the control and *gemin5^hg81^* mutant at 5 dpf after AG1478 treatment. Images were taken in the areas surrounding the end of yolk extension. White arrow points to the Tg(*SqET20*:EGFP) signal in the control embryo, which is dramatically increased in the mutant. Scale bar, 50 μm. The embryos used for the above analyses were generated from a pairwise incross of heterozygotic parents, treated with 2 μM AG1478 from 1 – 5 dpf, and then used Yopro-1 staining or transgenic fluorescence at 5 dpf to analyze neuromast formation.

### Neuromast cell proliferation is not induced by AG1478 in gemin5 mutants

To understand why *gemin5* mutants responded differently to AG1478, we used embryos with a double transgene Tg(*pou4f3*:GAP-GFP);(*SqET20*:EGFP) to label neuromast cells, and used an EdU incorporation assay to mark proliferating cells. We exposed the double transgenic embryos either to a mock treatment or to AG1478, and the resulting embryos were stained with the nuclear dye DAPI. In each neuromast, DAPI labeled all neuromast cells, the Tg(*pou4f3*:GAP-GFP) labeled hair cells, Tg(*SqET20*:EGFP) labeled mantle cells that demarcate the outer periphery of neuromast, and the GFP negative and DAPI positive cells in between were support cells. Quantification results showed that AG1478 promoted the proliferation of all three types of neuromast cells in wild-type larvae (Suppl. Fig. 5A-B), consistent with previous finding that AG1478 promotes cell proliferation (21).

We then applied AG1478 to *gemin5* embryos possessing the same Tg(*pou4f3*:GAP-GFP);(*SqET20*:EGFP) transgenes and monitored the neuromast cells in the *gemin5* mutant. When compared to the control siblings, *gemin5* mutants possessed a significantly reduced number of neuromast hair cells (as visualized by Tg(*pou4f3*:GAP-GFP) and mantle cells (as visualized by Tg(*SqET20*:GFP) (Fig. 4F), indicating that the *gemin5* mutation impaired neuromast cell proliferation in response to ErbB inhibition.

Several studies have indicated that AG1478 regulates neuromast cell proliferation through modulating the cell signaling activity between the Schwann cells, interneuromast cells, and the axons via WNT (21–24). We tested whether we could detect disruptions in Schwann cells in *gemin5* mutant embryos. Schwann cell morphology and quantity were evaluated using the Tg(*foxd3*:GFP) transgene or by the expression of *myelin basic protein a* (*mbpa*). Neither revealed a noticeable difference between the control siblings and mutants (Suppl. Fig. 6A-B). Lateral line axons were labeled with an antibody targeting acetylated tubulin and appeared to be comparable between the control and mutant embryos (Suppl. Fig. 6C). Wnt pathway activity was manipulated with both Wnt pathway activator BIO and inhibitor IWR1 and neither showed any differences between the mutant and control siblings (Suppl. Table 3). Altogether these data suggest the disruptions in myelination by the Schwann cells was not associated with the failure of AG1478 to induce supernumerary neuromasts in the *gemin5* mutant.

### Genetic mutations of ErbB pathway genes recapitulate the AG1478 effect

Similar to inhibition by AG1478, mutations in ErbB pathway genes, such as *erbb2, erbb3b* and *nrg1*, lead to an increase in lateral line neuromasts (21, 23, 24). Mutations in *erbb3b* and *nrg1* appear to have no other significant impact on embryo axis patterning nor on adult survival. We therefore generated stable genetic mutations for both *erbb3b* and *nrg1* and compared the effect of these mutations to the AG1478 effect on lateral line neuromast formation. As expected, both *erbb3b* and *nrg1* mutations caused a dramatic increase in the number of lateral line neuromasts, however, the increase was consistently lower than that of AG1478 treatment (Fig. 5A-B), suggesting AG1478 inhibits ErbB signaling more broadly than that mediated by either *erbb3b* or *nrg1* alone and that there may be some redundant signaling from other related genes.

**Fig. 5.**
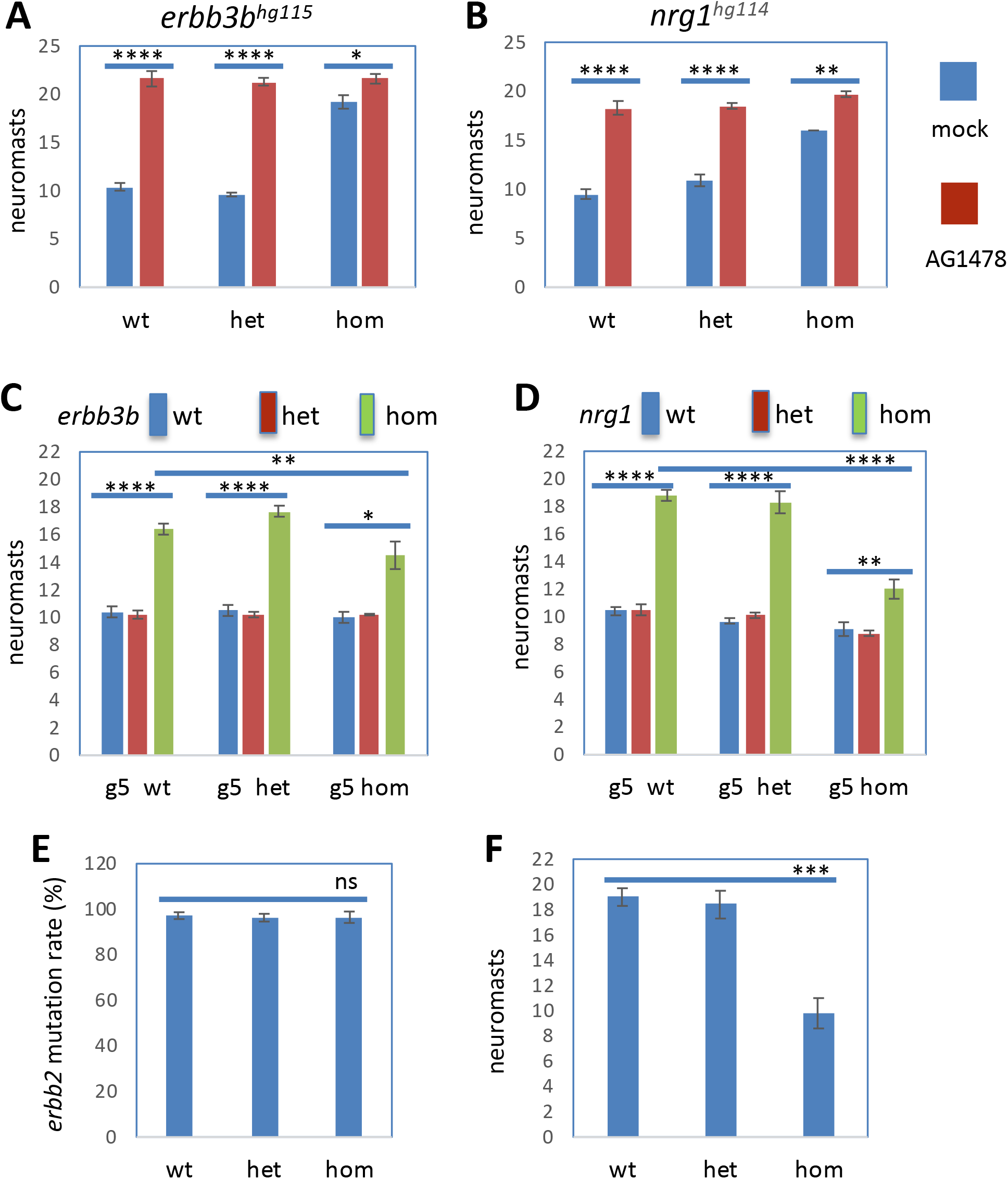
Reduced responsiveness of *gemin5^hg81^* mutants to the knockdown of ErbB pathway genes. (A) Quantification of lateral line neuromasts in the mock and AG1478 treated *erbb3b^hg115^* mutant embryos at 5 dpf. (B) Quantification of lateral line neuromasts in the mock and AG1478 treated *nrg1^hg114^* mutant embryos at 5 dpf. The embryos used for the *erbb3b^hg115^* and *nrg1^hg114^* mutation studies were generated from a pair of heterozygous parents, treated with 0 or 2 μM AG1478 from 1 – 5 dpf and then stained with Yopro-1 to count lateral line neuromasts. (C) Quantification of lateral line neuromasts in the *gemin5^hg81^/erbb3b^hg115^* mutant embryos at 5 dpf. The data are generated from analyzing 177 embryos generated from pairwise incrosses of double heterozygous parents and 5 embryos are double mutants. (D) Quantification of lateral line neuromasts in the *gemin5^hg81^/nrg1^hg114^* mutant embryos at 5 dpf. The data are generated from analyzing 156 embryos generated from pairwise incrosses of double heterozygous parents and 13 embryos are double mutants. (E) *erbb2* mutation rate in *erbb2* CRISPR guide RNA injected *gemin5* mutant embryos at 5 dpf. Mutation rate was measured by CRISPR-STAT fluorescent PCR-based fragment analysis (45). (F) Quantification of lateral line neuromasts in *erbb2* CRISPR guide RNA injected *gemin5* mutant embryos at 5 dpf. Error bars in the graphs indicate mean ± s.e.m. ns, P > 0.05; *P < 0.05; **P < 0.01; ****P < 0.0001.

We then examined how mutations of the genes in the ErbB pathway impact neuromast formation in the *gemin5* mutant background. We generated double mutants of *erbb3b/gemin5* or *nrg1/gemin5*. Consistent with our previous observations, a homozygous mutation in either *erbb3b* or *nrg1* alone caused an increase in the number of neuromasts, a homozygous mutation for *gemin5* alone caused no alteration, and a heterozygous mutation of either gene alone or together produced no change (Fig. 5C-D). Both double mutants displayed a lower level of increase of lateral line neuromasts when compared to *erbb3b* or *nrg1* mutant, however, the number of neuromasts in the double mutants was significantly higher when compared to the *gemin5* mutant alone, indicating disruption of the ErbB pathway could partially rescued the deficiency of neuromast formation in the mutant. The partial rescue in the double mutants suggest that AG1478 was failing to sufficiently inhibit ErbB signaling in *gemin5* mutants instead of the alternative explanation that interneuromast cells were unable to respond properly to release of ErbB signaling.

Rescue was also attempted by mutating the *erbb2* gene in the *gemin5* mutant background. Since *erbb2* loss of function is early embryonically lethal, we generated a mosaic knockdown of *erbb2* by injecting multiple CRISPR guide RNAs into the embryos from a *gemin5* heterozygous incross, and then used the injected embryos to quantify lateral line neuromast formation. Mutation frequency analysis showed these *erbb2* CRISPR guide RNAs resulted in a near-complete mutagenesis of the *erbb2* gene (Fig. 5E). Neuromast number quantification showed the *erbb2* knockdown promoted more neuromasts in the control sibling than in *gemin5* mutant (Fig. 5F).

### ErbB pathway inhibition partially rescues hair cell regeneration

Activation of ErbB signaling has been implicated in the regeneration of other tissues (25, 26), so we investigated whether ErbB pathway inhibition could improve hair cell regeneration in the three mutants that disrupt regeneration. In performing the hair cell regeneration assay in the presence of the ErbB inhibitor, AG1478 had no obvious effect on the regeneration of hair cells in control siblings, however, it did exhibit a dose-dependent rescue of regeneration in all three mutants (Fig. 6). Our interpretation of the data from Figures 5 and 6 is that ErbB signaling in the *smn1, gemin3*, and *gemin5* mutants was hyperactive, such that the increased ErbB activity was blocking AG1478 induction of ectopic neuromasts. Similarly, too much ErbB signaling was blocking the initiation of hair cell regeneration, but now AG1478 inhibition was sufficient to partially release the block in regeneration, presumably because ErbB signaling levels were generally lower in the regenerating neuromast compared to the interneuromast cells, or the level of reduction needed to see rescue was lower in the case of hair cell regeneration compared to neuromast induction.

**Fig. 6.**
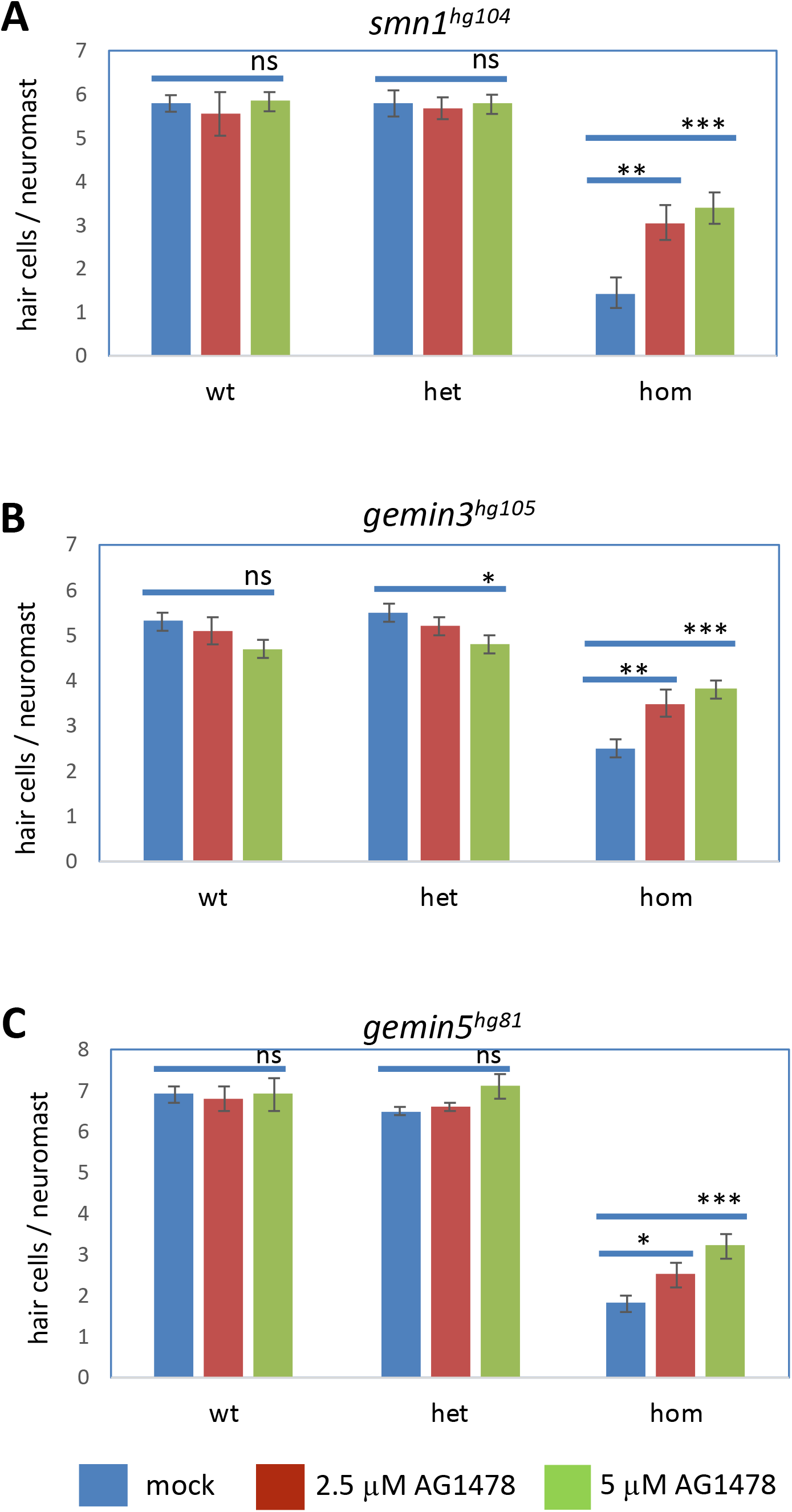
ErbB pathway inhibitor AG1478 partially rescues the hair cell regeneration deficiency in *gemin3, gemin5*, and *smn1* mutants. (A) Quantification of regenerated hair cells in the AG1478-treated *smn1^fh229^* mutant embryos at 2 days post hair cell ablation. (B) Quantification of regenerated hair cells in AG1478-treated *gemin3^hg105^* mutant embryos at 2 days post hair cell ablation. (C) Quantification of regenerated hair cells in lateral line neuromasts in AG1478-treated *gemin5^hg81^* mutant embryos at 2 days post hair cell ablation. The slight reduction in the hair cells of the *gemin3^hg105^* heterozygotes treated with 5 μM of AG1478 could be due to drug toxicity to this genetic background, since it was not observed in the *smn1^fh229^* and *gemin5^hg81^* embryos. Graphs show the mean ± s.e.m. Statistical difference are indicated as: ns, P > 0.05; *P < 0.05; **P < 0.01; ***P < 0.001. The embryos used for the analysis were generated from a single pair of heterozygous carrier parents, ablated hair cells at 5 dpf, and then treated with 0, 2.5 or 5 μM of AG1478 from 5 – 7 dpf.

### smn1, gemin3, and gemin5 mutations affect the regeneration of multiple tissues

Regeneration of different tissues can be achieved by utilizing a common set of molecular pathways and many genes are pan-regenerative in that they are induced and essential regardless of the specific injured tissue (17). Both *smn1* and *gemin5* genes were involved in regulating the regeneration of multiple tissues including neuromasts, caudal fins, and livers (10). To determine whether *gemin3* had similar phenotypes, we examined the regeneration of caudal fin and liver in *gemin3* mutants. Like in *smn1* and *gemin5* mutants, we found *gemin3* mutations did not alter the normal development of caudal fins or livers (data not shown), however, upon damage the mutant exhibited a deficiency in restoring the damaged tissues as was seen with the other mutants. After caudal fin amputation, the restored fin in the *gemin3* mutant was significantly smaller and often missing the pigment gap (Fig. 7A-B). Similarly, following chemical-mediated liver ablation in the Tg(*fabp10a*:CFP-NTR) transgenic background, the *gemin3* mutant displayed a clear deficiency in liver regeneration compared to the control siblings (Fig. 7C-D). As a control, *gemin6* mutants were also examined for a role in the regeneration of caudal fin and liver. In contrast to the regeneration mutants, *gemin6* mutants showed normal regeneration of both the caudal fin and the liver (Suppl. Fig. 7). These data suggest that *gemin3*, like *smn1* and *gemin5*, is generally involved in regeneration, regardless of the tissue tested.

**Fig. 7.**
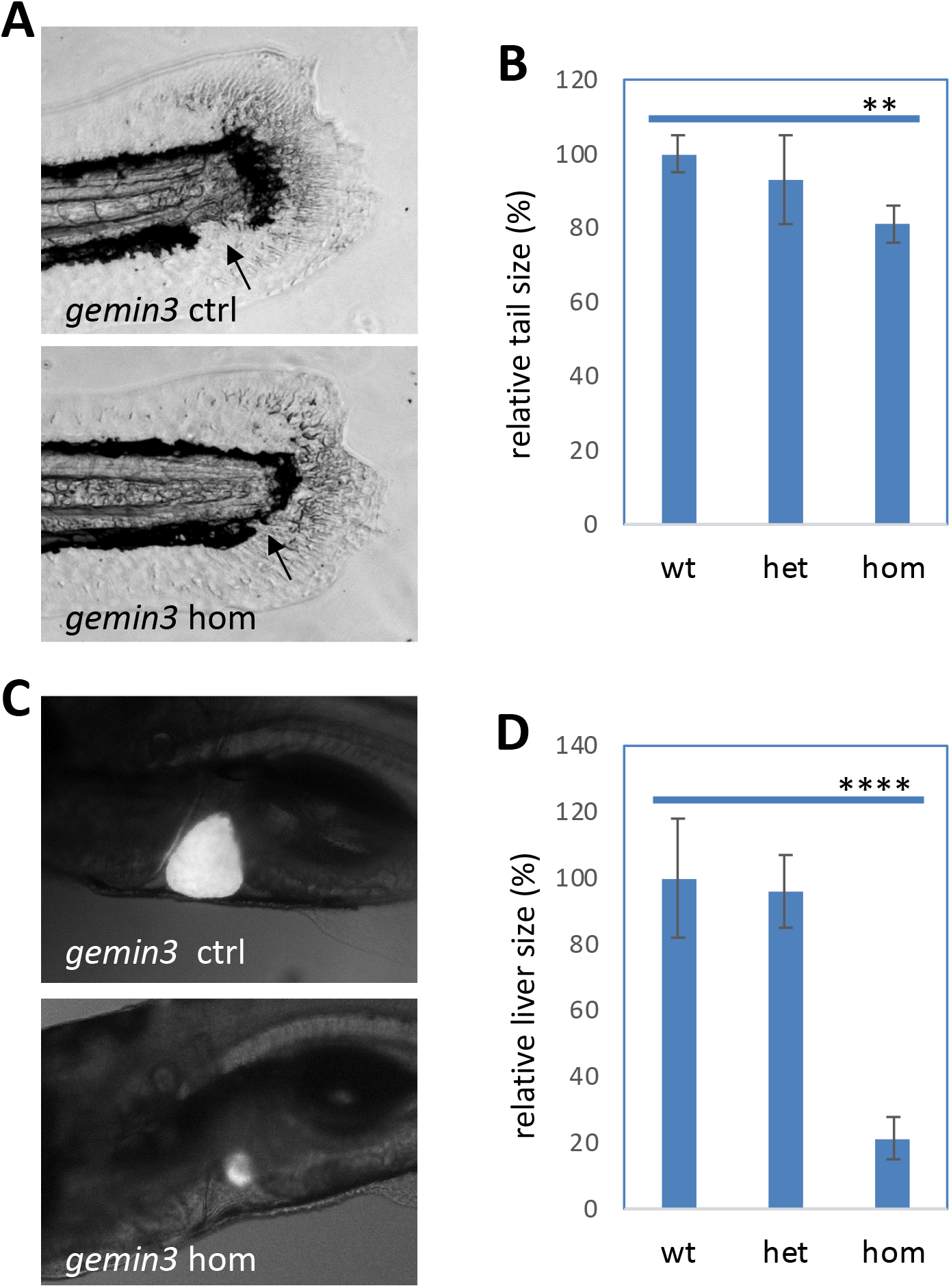
*gemin3^hg106^* mutations impair regeneration of caudal fins and livers. (A) Caudal fin regeneration in the control and *gemin3^hg106^* mutant embryos at 4 days post amputation. Arrows point to the pigment gap which is often missing in the mutants. (B) Quantification of the area of regenerated caudal fins in the *gemin3^hg106^* mutants. (C) Liver regeneration in the control and *gemin3^hg106^* mutant embryos at 3 days post ablation. (D) Quantification of the area of the regenerated livers. Representative images are shown. Liver tissue is labeled by Tg(*fabp10a*:CFP-NTR). Graphs show mean ± s.e.m. **P < 0.01; ****P < 0.0001.

### RNA-Seq reveals shared downstream targets among the genes involved in regeneration

To identify the pathways shared amongst the mutants blocking regeneration, we conducted RNA-sequencing (RNA-Seq) and miRNA-sequencing (miRNA-Seq) using the mutants of both regeneration genes and non-regeneration genes from the SMN complex. We found that the three genes impacting regeneration, regulated a common set of downstream targets which were distinct from the *gemin* mutations that did not affect regeneration (Fig. 8A-B). Significantly, we found *erbb3b* was upregulated in the three non-regenerative mutants (Fig. 8C), consistent with our hypothesis that ErbB signaling was hyperactive in these mutants (Fig. 5C). In addition, RNA-Seq data revealed that a mutation in one of the “regeneration genes” had no effect on the expression of the other two genes (Suppl. Fig. 8), suggesting there is no inter-regulation between the genes at the transcriptional level.

**Fig. 8.**
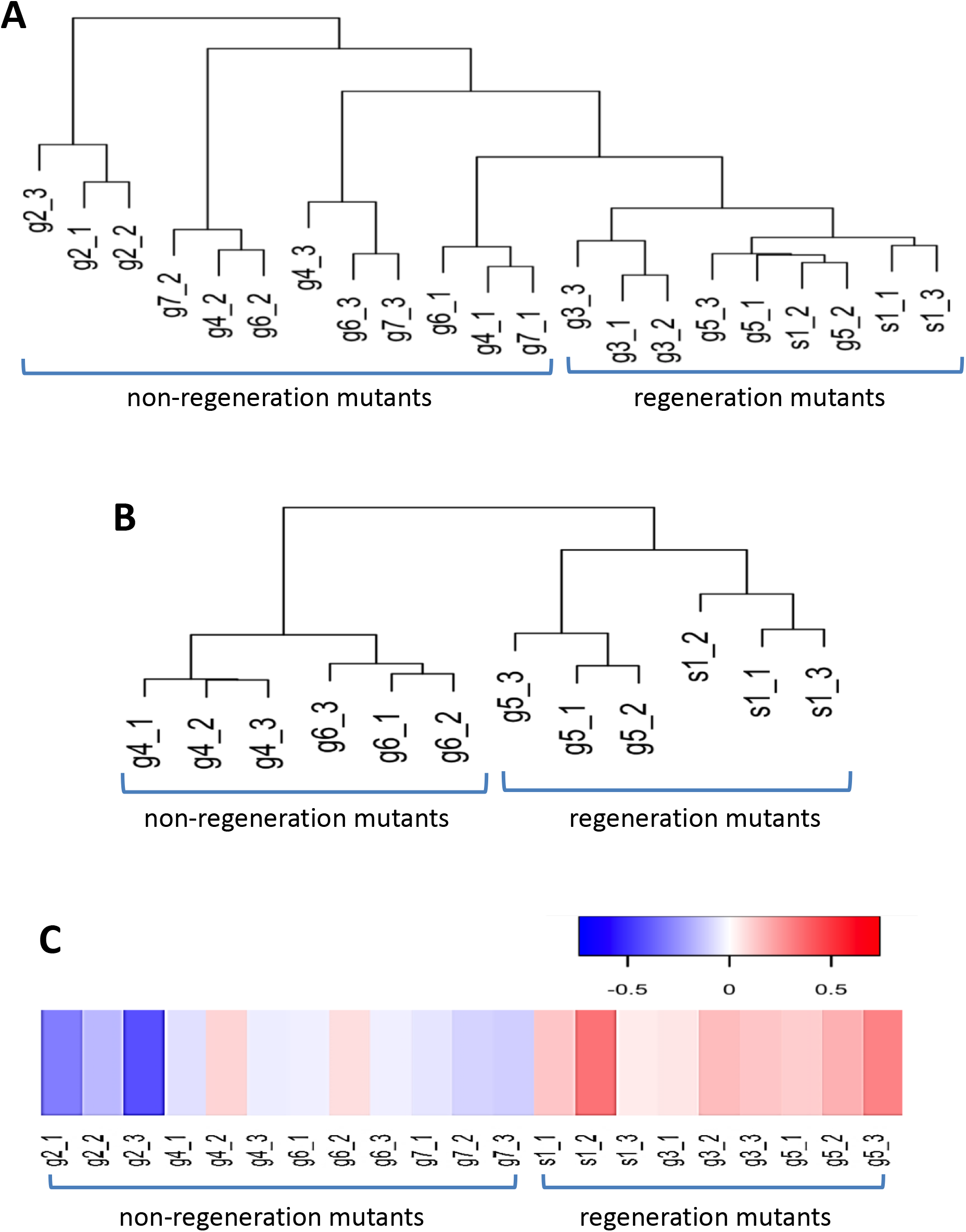
Shared genes dysregulated in *gemin3, gemin5*, and *smn1* mutants revealed by RNA-Seq and miRNA-Seq. (A) Hierarchical clustering of RNA-Seq samples using log2 fold change of normalized read counts. The control and mutant embryos used for the analysis were: *smn1^fh229^* (s1), *gemin2^hg108^* (g2), *gemin3^hg105^* (g3), *gemin4^hg109^* (g4), *gemin5^hg80^* (g5), *gemin6^hg110^* (g6) and *gemin7^hg111^* (g7). (B) Hierarchical clustering of miRNA-Seq samples using log2 fold change of normalized read counts. The control and mutant embryos used for the analysis were: *smn1^fh229^* (s1), *gemin4^hg109^* (g4), *gemin5^hg80^* (g5), and *gemin6^hg110^* (g6). The suffix numbers 1, 2 and 3 indicate the triplicates used for each sample. (C) Heat map of *erbb3b* expression in the RNA-Seq samples using log2 fold changes.

### Observed upregulation of the tp53/Mdm2 pathway was not the major cause of the regeneration phenotype

RNA-Seq data identified an increase in expression for both the *tp53* and *mdm2* genes specifically in the mutants inhibiting regeneration (Suppl. Fig. 9A). Several lines of published evidence indicate that p53 interacts with Mdm2 and activation of the p53/Mdm2 pathway is associated with SMN complex activity and SMA (27–29). To investigate a potential role for the tp53/Mdm2 pathway in hair cell regeneration, we depleted *tp53* genetically in both the *smn1* and *gemin5* mutant backgrounds. For the *gemin5* study, we crossed the *gemin5* mutant with the *tp53* M214K mutation (30), and found that the *gemin5/tp53* double mutants showed no improvement in regeneration or adult survival when compared to the *gemin5* mutant alone (Suppl. Fig. 9B, data not shown). For the *smn1* mutation, the *tp53* and *smn1* genes in zebrafish were both on chromosome 5, so we obtained double mutants carrying homozygous mutations for an *smn1* 2 bp insertion and a *tp53* 7 bp deletion, by injecting *smn1* CRISPR guide RNAs into embryos harboring a homozygous 7 bp *tp53* deletion mutation (31) and raised those fish for inbreeding. Consistent with the results of the *gemin5/tp53* double mutant, *tp53* mutants did not rescue the regeneration deficiency or the adult survival of *smn1* mutants (data not shown).

RNA-Seq data showed that Mdm2 was also significantly induced in the three genes involved in regeneration. Since *mdm2* mutations cause early embryonic lethality, we created a partial knockdown of Mdm2 by injecting *mdm2* CRISPR guide RNAs into the *gemin5* mutant background and found the resulting mosaic mutations in *mdm2* did not rescue the hair cell regeneration in the *gemin5* mutants (data not shown). Altogether, these data indicate that despite strong induction of *tp53* and *mdm2* in all three mutants blocking regeneration, the tp53/Mdm2 pathway is not a major contributor to the regeneration phenotype of the *gemin5* or *smn1* mutants although it does suggest that all three genes are involved in a common subset of pathways not shared by the other SMN complex proteins, and those pathways are involved in injury responses and tp53 stress responses.

## Discussion

Our previous large-scale mutagenesis screen showed *smn1* and *gemin5*, two SMN complex members, were essential for tissue regeneration (10). In this study, we expanded our mutagenesis screen to systemically examine the potential role of each of the nine members of the SMN complex in tissue regeneration. Consistent with the findings reported from other groups (32–34), we found that mutations in most SMN complex members were essential for adult survival (Suppl. Table 2). However, our genetic data suggest the nine SMN complex members can be categorized into three separate groups: *smn1, gemin3* and *gemin5* are required for both overall survival and regeneration after injury; *gemin2, gemin4, gemin6*, and *gemin7* are required for survival but not for regeneration; *gemin8* and *strap/unrip* appear to be non-essential factors for either regeneration or survival. The three regeneration members (*smn1, gemin3* and *gemin5*) are regulating regeneration through ErbB pathway-mediated cell proliferation, and they are essential for regeneration of multiple (if not all) tissues.

Studies of the SMN complex has been ongoing for more than two decades (35) with the largest focus on *SMN1* because mutations in this gene are responsible for the human disease spinal muscular atrophy (1, 2). However, the association of SMN complex members with tissue regeneration was not recognized until our prior study (10) and expanded here. Our work strongly suggests that some of the SMN complex members have separate, independent functions unrelated to snRNP assembly, or that the complex isn’t always functioning with all nine genes in a stable, stochiometric ratio. For example, we found transcripts were not expressed uniformly and ubiquitously, but expression varied in different brain regions and in different neuromast cells (Suppl. Fig. 2). However, no additive or synergistic interactions were observed between the three genes involved in regeneration (Suppl. Fig. 3). All three regeneration members functioned through ErbB-pathway-mediated cell proliferation (Fig. 4 and 6), all three possessed an ability to regulate regeneration in multiple tissues (Fig. 7) (10), and none of the three appeared to be epistatic to the other two. All these findings argue that these regenerative members work together in a shared molecular mechanism. Our findings suggest that the three SMN complex members involved in regeneration possess functions separate from snRNP biosynthesis that are essential for tissue regeneration and are also related to tp53 regulation/activation, although our genetic evidence in these two functions are not directly related.

In line with our findings, previous studies have reported apparently independent activities of SMN complex members. For example, SMN1 has been implicated in motor neuron growth (36, 37), SMN1’s function in motor neurons appears to be independent of snRNP biosynthesis (38), and SMN1 has a specific role in axonal mRNA regulation and axonogenesis (5, 6). Furthermore, GEMIN3, an RNA helicase, is involved in cell proliferation and microRNA regulation of signal transduction (7). Gemin5 regulates *smn1* expression (20), and Gemin5’s C-terminus can regulate protein synthesis (8, 39). Future studies should be able to evaluate SMN complex-dependent and independent functions more precisely through detailed analysis of splicing isoforms in different genes and/or cell types, under natural, diseased or regenerative conditions.

In this study, we demonstrated a link between the ErbB pathway and three of the SMN complex’s proteins. Chemical inhibition of the ErbB pathway with AG1748 in *smn1, gemin3*, and *gemin5* mutants failed to induce ectopic neuromasts or rescue hair cell regeneration, while genetic ablation of *erbb3b* or *nrg1* was able to partially rescue the neuromast induction and hair cell regeneration, suggesting the ErbB pathway is hyperactive in the mutants. Because the ErbB pathway is associated with various neurological diseases (40), it suggests future investigation is warranted to address whether the upregulation of the ErbB pathway in the three SMN mutants is specific to injury responses or if it is also one of the underlying mechanisms in the neurodegenerative pathology of SMA.

Several studies have demonstrated that the ErbB pathway plays a promotive role in the regeneration of other tissues. For example, mutations in *erbb2* or *erbb3* cause a deficiency in caudal fin regeneration (26), and AG1478 treatment inhibits the regenerative proliferation of cardiomyocytes (25). Our data indicate that the role of the ErbB pathway in regeneration differs based on tissue type. It remains unclear how the ErbB signaling is integrated into the different roles it plays in different tissues.

Our RNA-Seq data revealed *erbb3b* is upregulated in *smn1, gemin3*, and *gemin5* mutants (Fig. 8C) and we found that inhibition of ErbB pathway contributes to a partial rescue of their regeneration phenotype (Fig. 6), suggesting that the ErbB pathway is, at least in part, the underlying mechanism for the deficient regeneration, and likely it is only one of many pathways affected during the regeneration. Besides *erbb3b, p53* and *mdm2* were also upregulated (Suppl. Fig. 9A). The p53/Mdm2 pathway has long been documented to interact with the SMN complex. p53 has a direct physical interaction with both SMN1 and Gemin3 (29, 41). p53 depletion rescues *mdm2* mutant phenotypes (27). Abnormal *mdm2* splicing and p53 activation are associated with the death of motor neurons in SMA (28). We found inhibition of the p53/Mdm2 pathway brought no alteration to survival or regeneration in the *gemin5* mutants (Suppl. Fig. 9. Data not shown), suggesting this pathway is not the major cause of the mutant regeneration phenotypes.

In conclusion, this study provides insight into the SMN complex and potential roles for the complex in wound healing and ErbB signaling. Although *SMN1* is the causative gene in the majority of SMA patients, there are still cases of SMA where the causative gene is unknown. Because we see phenotypes cluster with *smn1, gemin3*, and *gemin5*, it is possible that a fraction of undiagnosed SMA cases or related neurodegenerative diseases could be caused by variants in either *GEMIN3* or *GEMIN5*. It is also possible that the functions of the three SMN complex members outside of snRNP assembly are somehow linked to SMA pathology and deficient regeneration is an underlying mechanism for SMA and even for other neurological diseases.

## Materials and Methods

### Zebrafish husbandry and embryology

Zebrafish husbandry and embryo staging were performed according to Kimmel (42). All experiments were in compliance with NIH guidelines for animal handling and research and approved by the NHGRI Animal care and Use Committee (protocol G-01-3). Adult fish survival was examined at 3 months post fertilization. Quantitative PCR (qPCR) was conducted by extracting total RNA with Trizol (Invitrogen, Cat#: 15596026), synthesizing cDNA with SuperScript first-strand synthesis system (Thermo Fisher Scientific. Cat#: 11904018), and then running qPCR with SYBR^™^ Green PCR Master Mix (Thermo Fisher Scientific, Cat#: 4344463). Beta-actin was used as an internal reference. Semi-qPCR analysis was conducted similarly as qPCR excepted no use of SYBR Green and amplicons analyzed on an agarose gel. CRISPR/Cas9 mutagenesis was performed as previously described (43). The CRISPR targets and primers used for mutation detection are listed in the CRISPRz database (44) https://research.nhgri.nih.gov/CRISPRz/). CRISPR mutation rates for founder embryos were analyzed by calculating the percentage of mutant signal over the total signal (45).

### Biological materials and the zebrafish transgenic lines

The biological dyes used in this study were: Yopro-1 (Life Technologies. Cat#: Y3603), ProLong Gold Antifade Mountant with DAPI (Vector Laboratories. Cat#: H1200). Chemical compounds used in this study were all purchased from Sigma: copper sulfate (C1297), antimycin A (A8674), cycloheximide (C7698), AG1478 (T4182), DAPT (D5942), dexamethasone (D4902), prednisolone (P6004), 1-azakenpaullone (A3734), BIO (B1686), IWR1 (I0161), SU5402 (SML0443), SB505124 (S4696), H_2_O_2_ (216763), NAC (A7250), and GSH (G4251). All chemicals except NAC and GSH were dissolved in DMSO. NAC and GSH were dissolved in embryo media 1x Holtfreter’s buffer. Chemical treatments were carried out in embryo media, with the doses and durations listed in Suppl. Table 3. Mock treatments were carried out by adding an equal amount of the corresponding solvents. The zebrafish transgenic lines used were: Tg(*atoh1a*:dTomato)^nns8^ (46), Tg(*tnks1bp1*:EGFP) (47), Tg(*clndb*:GFP)^zf106^ (48), Tg(*pou4f3*:GAP-GFP)^s273t^ (49). Tg(*SqET20*:EGFP) (50), Tg(*foxd3*:GFP)^zf15^ (51), Tg(*fabp10a*:CFP-NTR)^s931^ (52). Imaging of the transgenic embryos were conducted using either an inverted Zeiss Axiophot or a Zeiss LSM 880 confocal microscope.

### Hair cell and neuromast quantification

Hair cell staining and quantification were as described (53). Briefly, for analyzing hair cell development, embryos from heterozygotic incrosses were cultured until 5 dpf, and then placed in a cell strainer (BD Falcon) for staining with 2 μM YoPro-1 for 30 minutes. Lateral line neuromasts P1, P2, P4 and P5 in each embryo were used for hair cell counting using an inverted Zeiss Axiophot. The number of neuromasts in the lateral line of each embryo were also counted for studying neuromast formation. For hair cell regeneration analysis, embryos from heterozygotic incrosses at 5 dpf were treated with the ototoxin copper sulfate at 10 μM for 2 hours except when otherwise indicated, recovered for 48 hours, and then counted for the regenerated hair cells in the lateral line neuromasts P1, P2, P4 and P5. Approximately 40 embryos were used for each of the analyses except when otherwise indicated. The average number of hair cells and the standard error of the mean (s.e.m.) are shown in the graphs.

### Immunohistochemical staining

Hair cell staining for fixed zebrafish tissues were performed with a combination of antibodies against hair cell soma-1 (Developmental Studies Hybridoma Bank. Cat#: HCS-1, 1 μg/ml) and myosin-VIIa (Developmental Studies Hybridoma Bank. Cat#: 138-1, 1 μg/ml), followed by an Alexa 488-labeled secondary antibody (Invitrogen. Cat#: A11001, 4 μg/ml). Alkaline phosphatase staining for lateral line neuromasts were performed as previously reported (21). Lateral line axons were stained with an antibody against acetylated tubulin (Sigma. Cat#: T7451, 1:1000 dilution) and a secondary antibody conjugated with Alexa 594 (Invitrogen. Cat#: A11012, 4 μg/ml). Proliferating cells were labeled with the Click-It EdU Alexa Fluor 594 imaging Kit (Life Science. Cat#: C10339), following the manufacturer’s instructions. Embryos were prepared by exposure to 500 μM of EdU in 1x Holtfreter’s buffer with 15% DMSO in an ice bath for 30 minutes, recovered for 3 hours, and then fixed in 4% paraformaldehyde overnight. The stained embryos were mounted with ProLong Gold Antifade Mountant with DAPI on a microscope slide and imaged with a Zeiss LSM 880 confocal microscope. The neuromast images were then used for EdU positive cell quantification. For the *smn1* mutation, neuromasts were additionally labeled by Tg(*cldnb*:GFP). For the *gemin5* mutation, neuromasts were additionally labeled by Tg (*tnks1bp1*:EGFP).

### Quantifying development and regeneration of caudal fin

Caudal fin development and regeneration were analyzed as previously described (10). In brief, embryos were obtained from a pair of heterozygous parents. Fin development was measured at 5 dpf, using the posterior of pigment gap as a positional reference. For the regeneration analysis, amputation was performed at 3 dpf, at the posterior end of ventral pigment break. The regeneration was measured at 7 dpf, continuing to use the anterior end of pigment gap as a positional reference. ImageJ was used for quantifying the fin areas. All analyzed embryos were genotyped. Graph shows the mean and s. e. m, based the quantification data from approximately 10 embryos per genotype.

### Quantification of development and regeneration of liver

Liver development and regeneration was tested using the transgene Tg(*fabp10a*:CFP-NTR (52). The embryos used for the analysis were the CFP-positive embryos obtained from a pair of parents with one carrying the heterozygous gene mutation and the other carrying both the heterozygous mutation and an allele of Tg(*fabp10a*:CFP-NTR). Liver size was measured at 5 dpf. For liver regeneration analysis, the embryos were treated with 10 mM metronidazole for 1.5 days at 3 dpf and analyzed for regeneration at 7 dpf. All analyzed embryos were imaged at a lateral view with head facing right under a Zeiss Axiophot fluorescent microscope, and afterwards genotyped. ImageJ was used to measure the liver areas. Approximately 45 CFP-positive embryos were used for each analysis. Graph shows the mean and s. e. m.

### RNA-Seq and miRNA-Seq analyses

The embryos used for RNA-sequencing (RNA-Seq) and miRNA-sequencing (miRNA-Seq) were produced from a cross of a single pair of heterozygous parents, exposed to 10 μM copper sulfate for 2 hours at 5 dpf, and then subjected to caudal fin biopsy for genotyping and the body stored in Qiazol (Qiagen. Cat#: 79306) at 7 dpf. Afterwards the wild-type and homozygous mutant embryos were pooled together and used for total RNA extraction by using Qiagen miRNeasy Mini Kit (Cat#: 217004). The total RNA with an integrity score (RIN) over 9 were used for RNA-Seq and miRNA-Seq analyses.

### Statistical analyses

A student t-test (two tailed) was used for comparison between two samples. One-way ANOVA was used when comparing multiple samples. A difference was considered significant when P value was less than 0.05. Error bars in the graphs represent mean ± s.e.m. Asterisks and short lines were used to indicate a significant difference between two groups. ns, P >= 0.05; *P < 0.05; **P < 0.01; ***P < 0.001; ****P < 0.0001. Each experiment presented was repeated at least twice, with the replicates showing statistical significance each time.

## Acknowledgement

We would like to thank MaryPat Jones, Blake Carrington, Kevin Bishop and Raman Sood from the NHGRI Zebrafish Core for mutation genotyping; Suiyuan Zhang from the NHGRI Bioinformatics Core for CRISPRz database management; Alisha Beirl and Katie Kindt from the NIDCD for mechanistic study; Daniel Green, Justin Frye, Jason Frye, Hillary Mahon and Paulina Capar in Charles River for fish care; and the members of Burgess laboratory for helpful discussion. We have complied with all relevant ethical regulations regarding animal use and all animal experiments were approved by the National Human Genome Research Institute’s Animal Care and Use Committee (protocol #G-01-3). This research was supported by the Intramural Research Program of the National Human Genome Research Institute (ZIAHG200386-06).

## Supplemental Figure Legends

**Suppl. Fig. 1.**
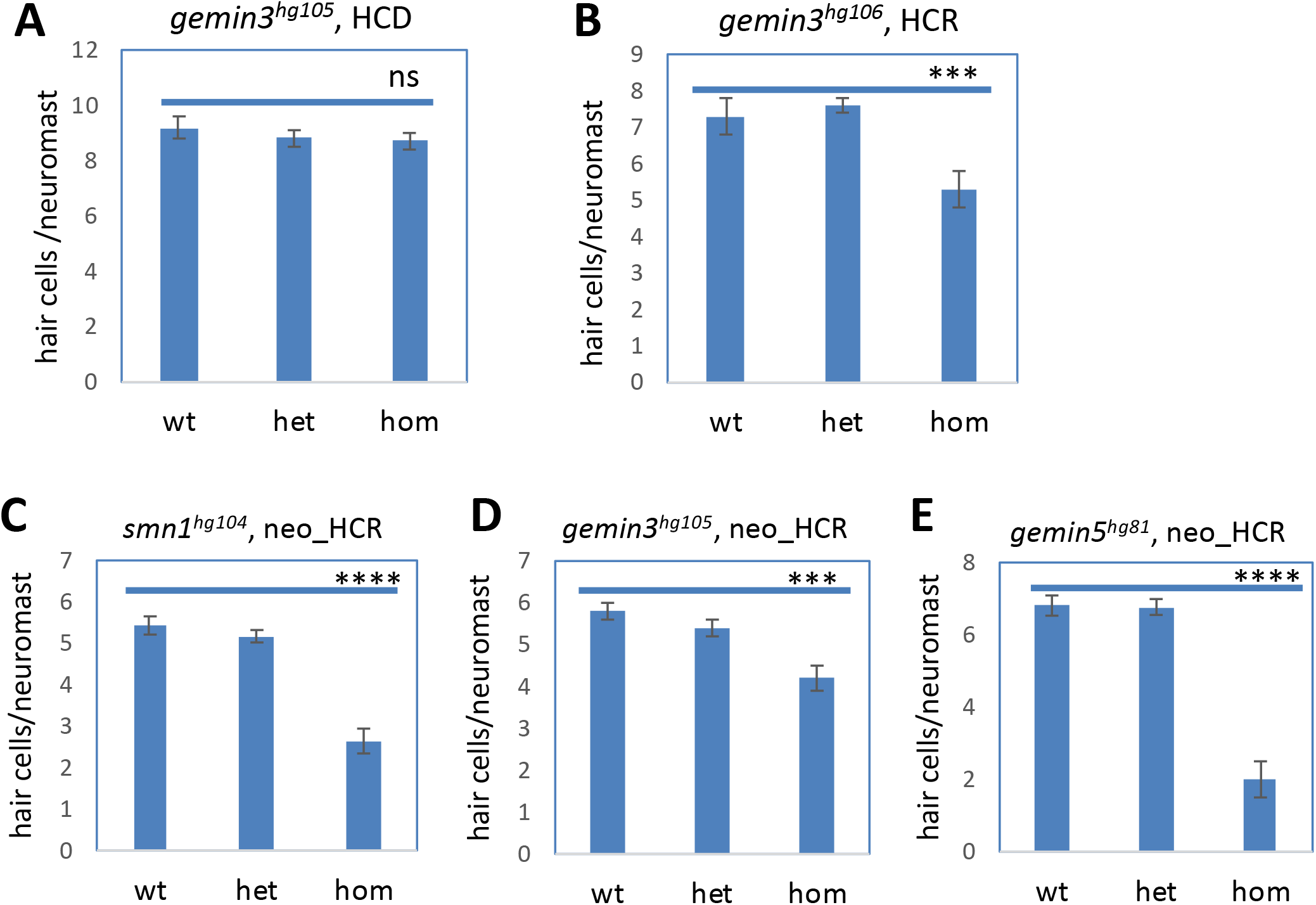
Hair cell development and regeneration in *gemin3, gemin5*, and *smn1* mutants. (A) Normal hair cell development in *gemin3^hg105^* mutants. (B) Impaired hair cell regeneration in *gemin3^hg106^* mutants. (C-E) impaired hair cell regeneration after neomycin-induced hair cell ablation in homozygous mutations of *smn1^hg104^* (C), *gemin3^hg105^* (D) and *gemin5^hg81^* (E). HCD, hair cell development. HCR, hair cell regeneration. neo, neomycin. wt, wild-type. het, heterozygotes. hom, homozygotes. Error bars in the graphs represent mean ± s.e.m. ns, P > 0.05; ***P < 0.001; ****P < 0.0001.

**Suppl. Fig. 2.**
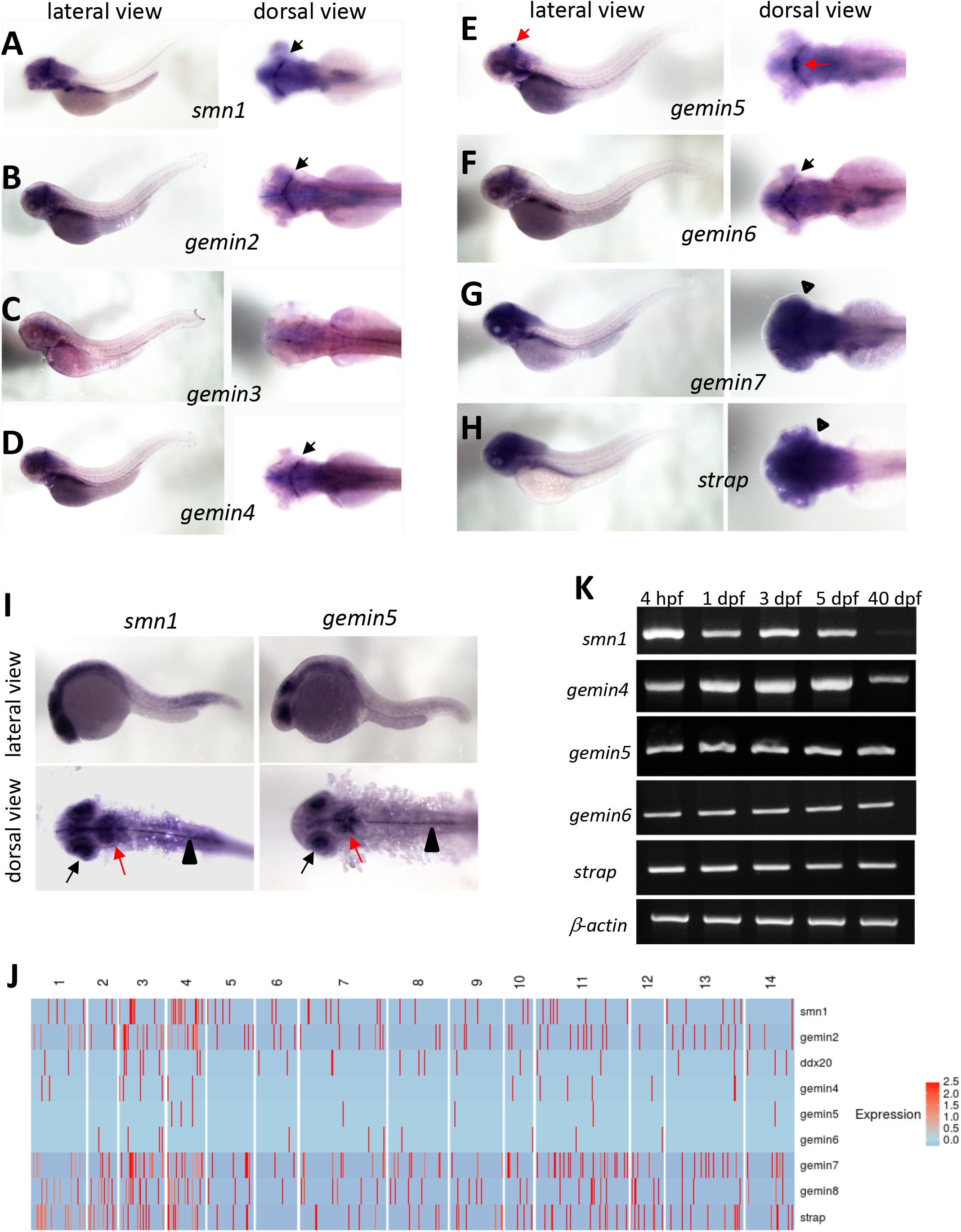
Expression analysis of the SMN complex genes. (A-H) Whole-mount in situ hybridization analysis using TAB5 wild-type embryos at 3 dpf to detect the expression patterns of *smn1* (A), *gemin2* (B), *gemin3* (C), *gemin4* (D), *gemin5* (E), *gemin6* (F), *gemin7* (G) and *strap* (H). Black arrows point to the restricted brain expression of *smn1, gemin2, gemin4*, and *gemin6*. Red arrows point to the unique expression of *gemin5* as a stripe with a thickening in the midline. Black arrowheads point to the ubiquitous expression of *gemin7* and *strap*. No obvious expression of *gemin8* was detected at this embryonic stage. (I) Whole-mount in situ hybridization analysis using TAB5-wild type embryos at 1 dpf to detect the expression patterns of *smn1* and *gemin5*. For imaging the dorsal view, yolks of the embryos were removed for better visualization of the staining. Black arrows point to expression in the eye. Red arrows point to the expression in the forebrain. Black arrowheads point to midline expression. (J) Heatmap of expression of the SMN complex genes in the scRNA-sequencing data reported by Lush ME, et al (13). Each red hash indicates a cell that expresses the specific gene. The numbers 1-14 indicate the different clusters of neuromast cells, with clusters 1 and 2 indicating hair cells, clusters 5 and 6 indicating mantle cells, and the others indicating support cells. The *gemin5* gene is not expressed in hair cells (clusters 1 and 2). The expression of *ddx20/gemin3, gemin4, gemin5* and *gemin6* are overall weaker than the other genes. (K) Semi-quantitative PCR analysis of the expression levels of SMN complex genes at different developmental stages.

**Suppl. Fig. 3.**
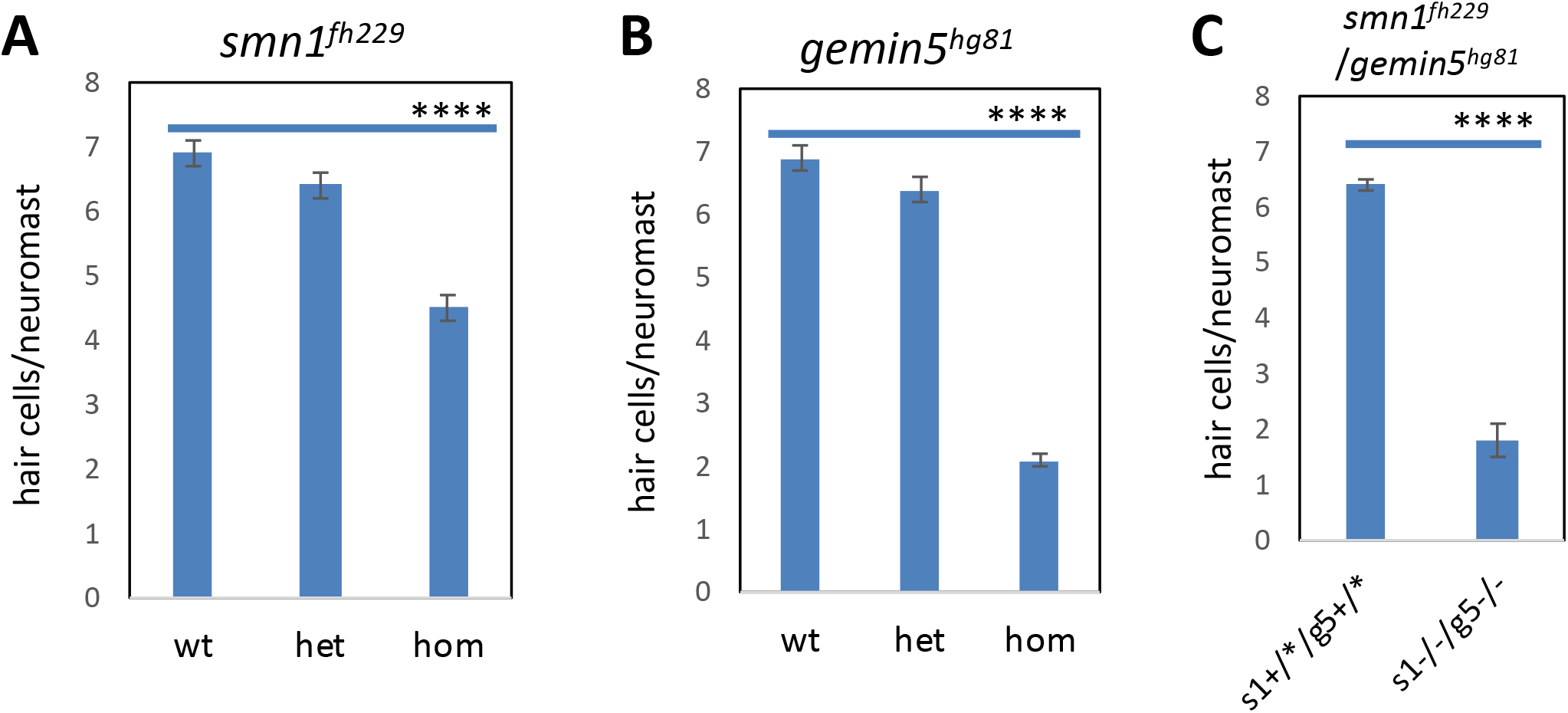
Hair cell regeneration in *smn1^fh229^* and *gemin5^hg81^* single and double mutants. The embryos used for the analysis were generated from a single pair of parents, each carrying heterozygous mutations for both *smn1^fh229^* and *gemin5^hg81^*. Graphs show the hair cells regenerated from the *smn1^fh229^* mutant (A), *gemin5^hg81^* mutant (B), and *smn1^fh229^/gemin5^hg81^* double mutant (C). The difference is significant between the *smn1* wild-type and homozygotes, between the *gemin5* wild-type and homozygotes, and between the *smn1/gemin5* control and double mutant (**** P < 0.0001 for all three groups). There is no difference between the *gemin5* homozygotes and the *smn1/gemin5* double homozygotes (ns, P > 0.05. Not labeled in the graphs).

**Suppl. Fig. 4.**
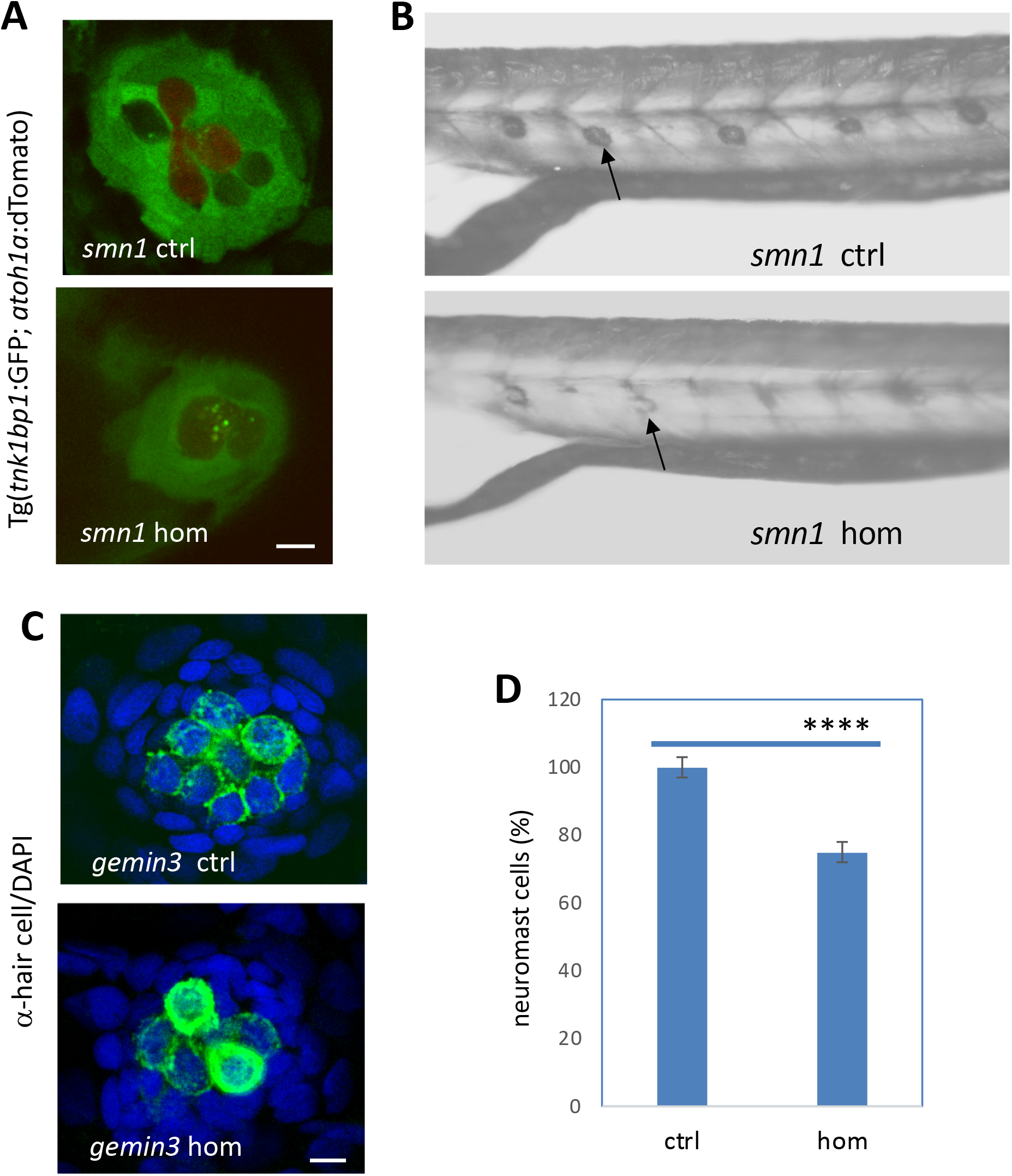
Smaller neuromasts observed in *smn1^fh229^* and *gemin3^hg105^* mutants at 2 days post hair cell ablation. (A) Imaging the lateral line neuromasts in the *smn1^fh229^* control and mutant embryos at 2 days post hair cell ablation, using Tg(*tnks1bp1*:GFP) and Tg(*atoh1a*:dTomato). Scale bar, 10 μm. (B) Alkaline phosphatase staining of lateral line neuromasts in the *smn1^fh229^* control and mutant embryos at 2 days post hair cell ablation. Arrows point to the neuromasts. (C) Confocal images of lateral line neuromasts in the *gemin3^hg105^* control and mutant embryos at 2 days post hair cell ablation. Neuromasts were stained with anti-hair cell antibodies (green) and DAPI (blue). Scale bar, 10 μm. (D) Quantification of neuromast cells in the *gemin3^hg105^* control and mutant embryos. Error bars in the graphs represent mean ± s.e.m. A significant reduction is found in the neuromast cells (**** P < 0.0001). The numbers are presented as percentages based on cell numbers in confocal images.

**Suppl. Fig. 5.**
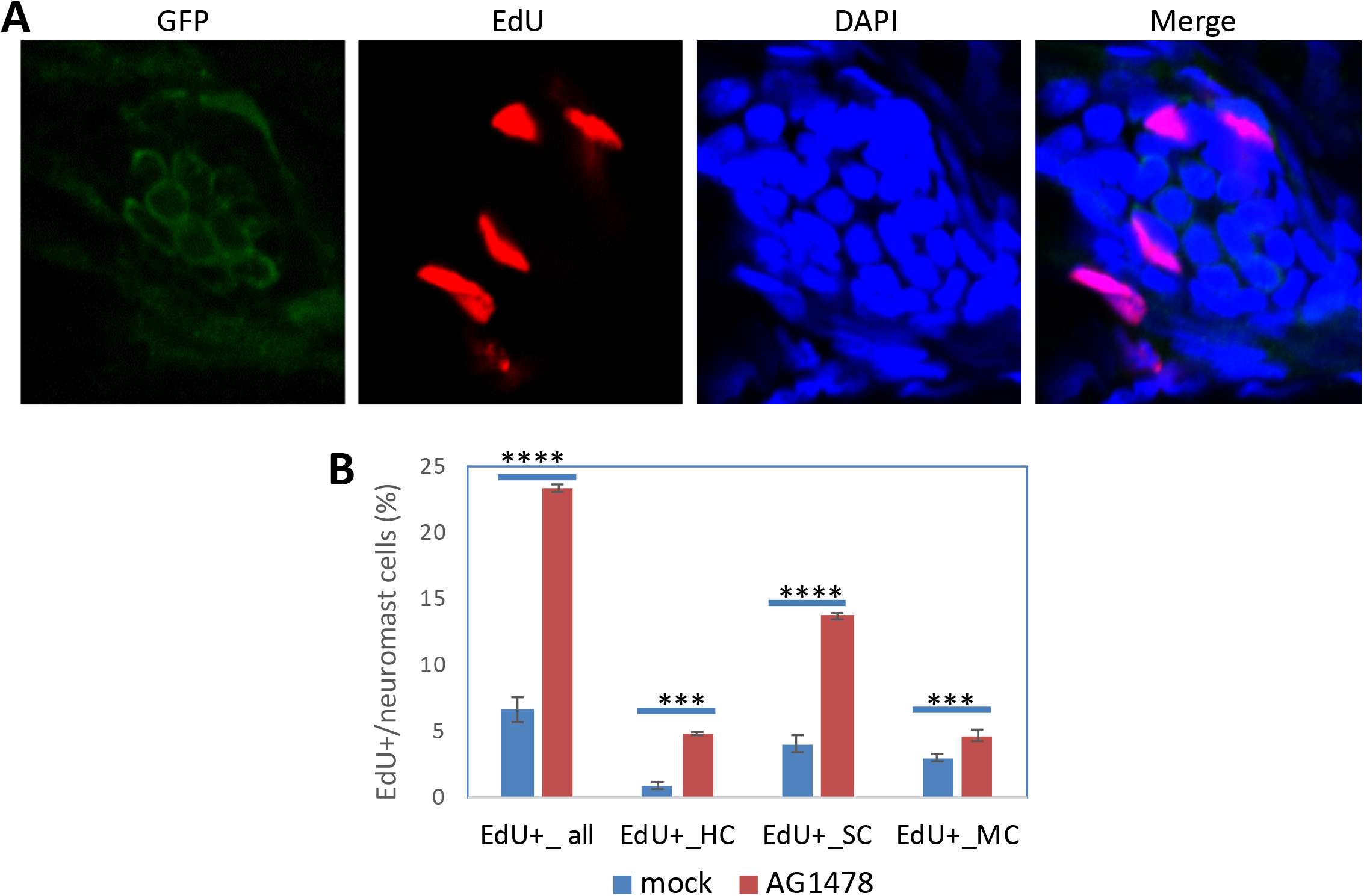
ErbB pathway inhibitor AG1478 promotes lateral line neuromast cell proliferation. (A) Confocal images of a lateral line neuromast of a TAB5 embryo at 5 dpf labeled by transgenic GFP from Tg(*pou4f3*:GAP-GFP) and Tg(*SqET20:EGFP*), EdU, DAPI and the merged. Representative images are shown. Scale bar, 10 μm. The embryos were treated with 2 μM AG1478 from 1 – 4 dpf and then used for EdU labeling. (B) Quantification of the proliferating cells in neuromasts (EdU+_all), hair cells (EdU+_HC), support cells (EdU+_SC), and mantle cells (EdU+_MC) in the mock and AG1478-treated TAB5 embryos at 5 dpf. Error bars represent mean ± s.e.m. Statistical difference is labeled as ***P < 0.001 and ****P < 0.0001.

**Suppl. Fig. 6.**
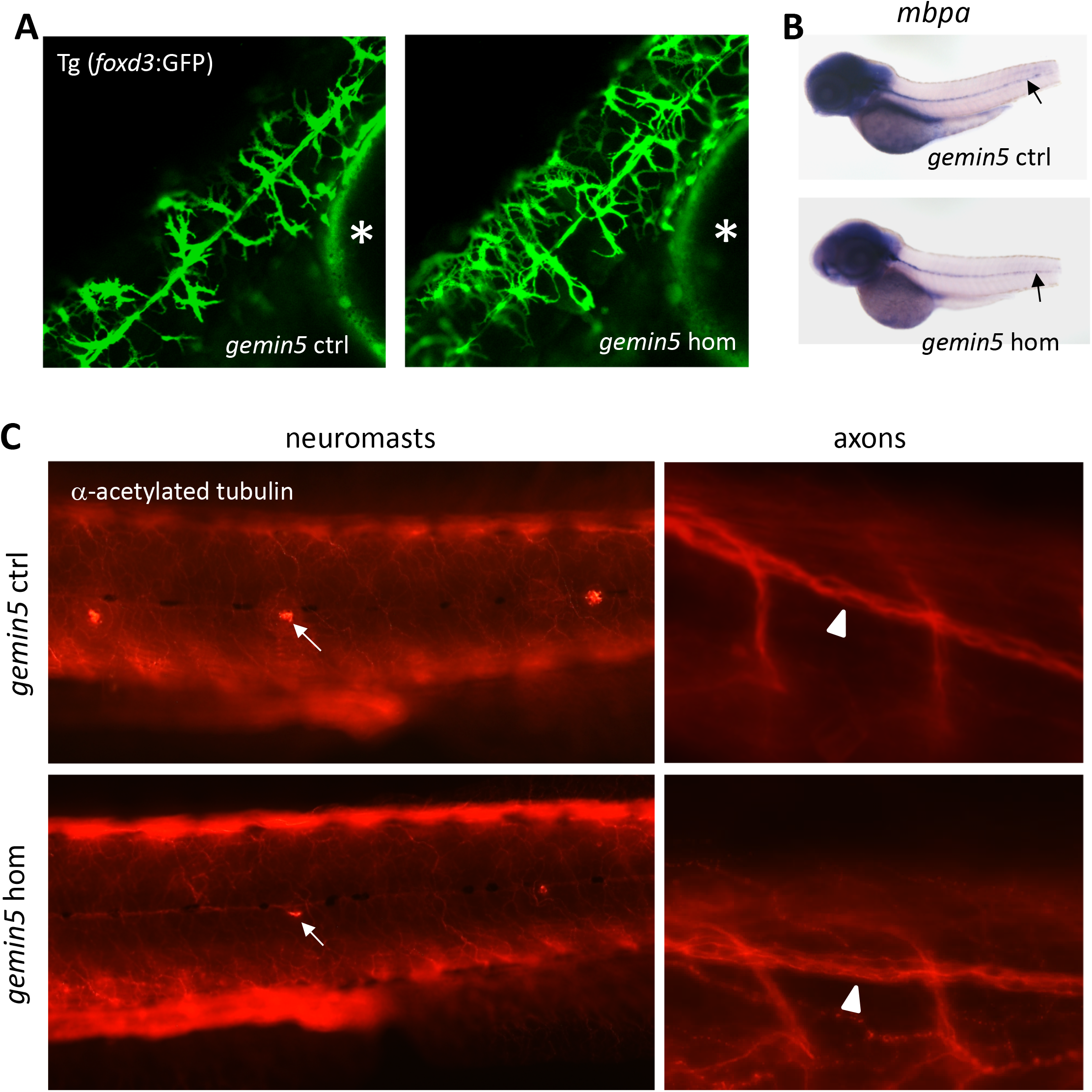
Normal development of Schwann cells and lateral line axons in the *gemin5^hg81^* mutant. (A) Confocal images of Schwann cells in the lateral line of the control and *gemin5^hg81^* mutant at 2 dpf. Asterisks label the embryonic yolk. (B) Whole-mount in situ hybridization analysis of *myelin basic protein a* (*mbpa*) expression in the control and *gemin5^hg81^* mutant at 4 dpf. Arrows point to the expression in the lateral line. The caudal fin folds were amputated to obtain tissue for mutation genotyping. (C) Fluorescent images of the lateral line axons labeled by anti-acetylated-tubulin antibodies in the control and *gemin5^hg81^* at 2 days post hair cell ablation. Representative images are shown. Arrows point to lateral line neuromasts. Arrowheads point to the lateral line axons.

**Suppl. Fig. 7.**
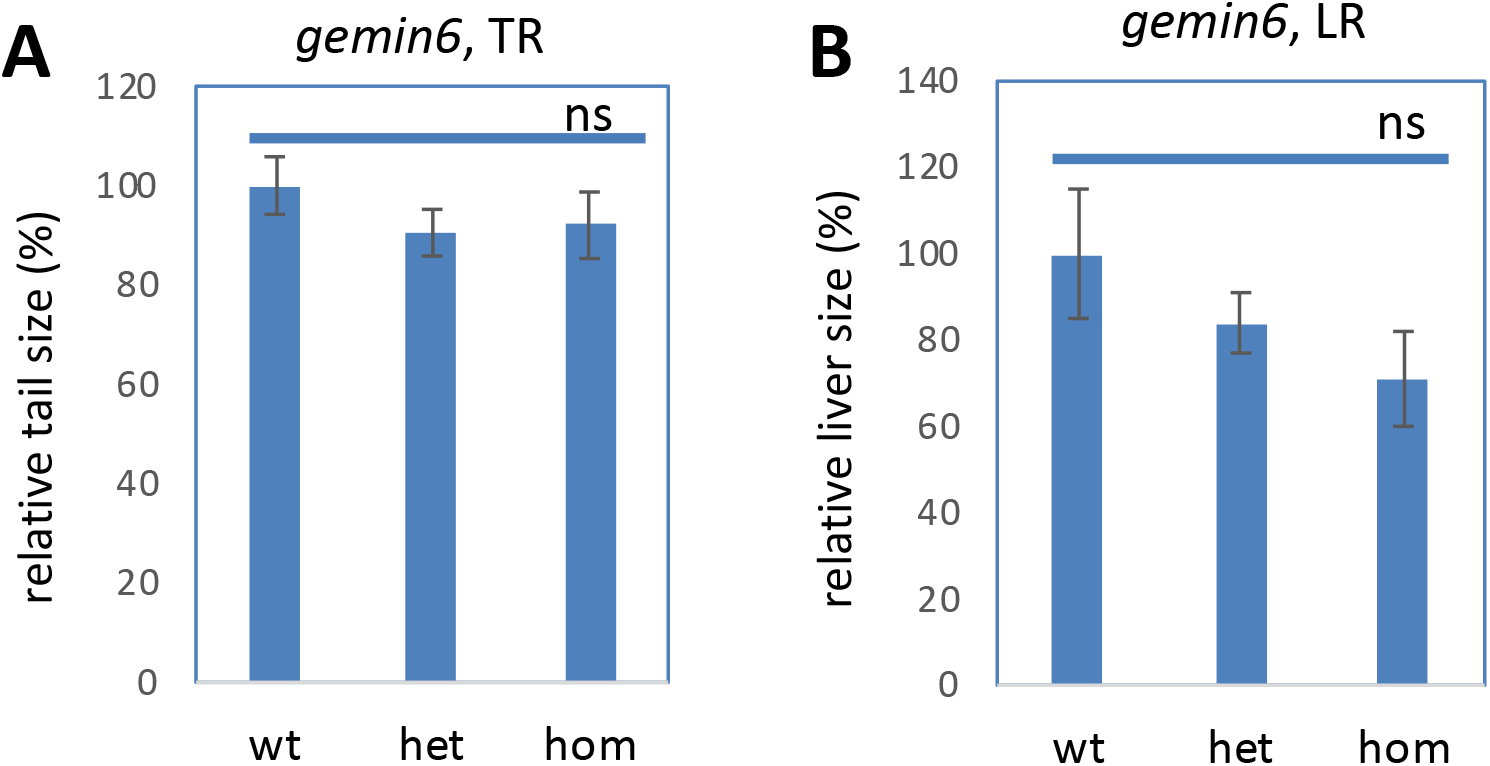
The *gemin6^hg110^* mutation had no impact on the regeneration of caudal fin folds or livers. (A) Quantification of the area of the regenerated caudal fins. (B) Quantification of the area of the regenerated livers. Liver is labeled by Tg(fabp10a:CFP-NTR). Graphs show the mean ± s.e.m. The difference between the wild-type (wt) and homozygous mutants (hom) were not significant (ns, P > 0.05).

**Suppl. Fig. 8.**
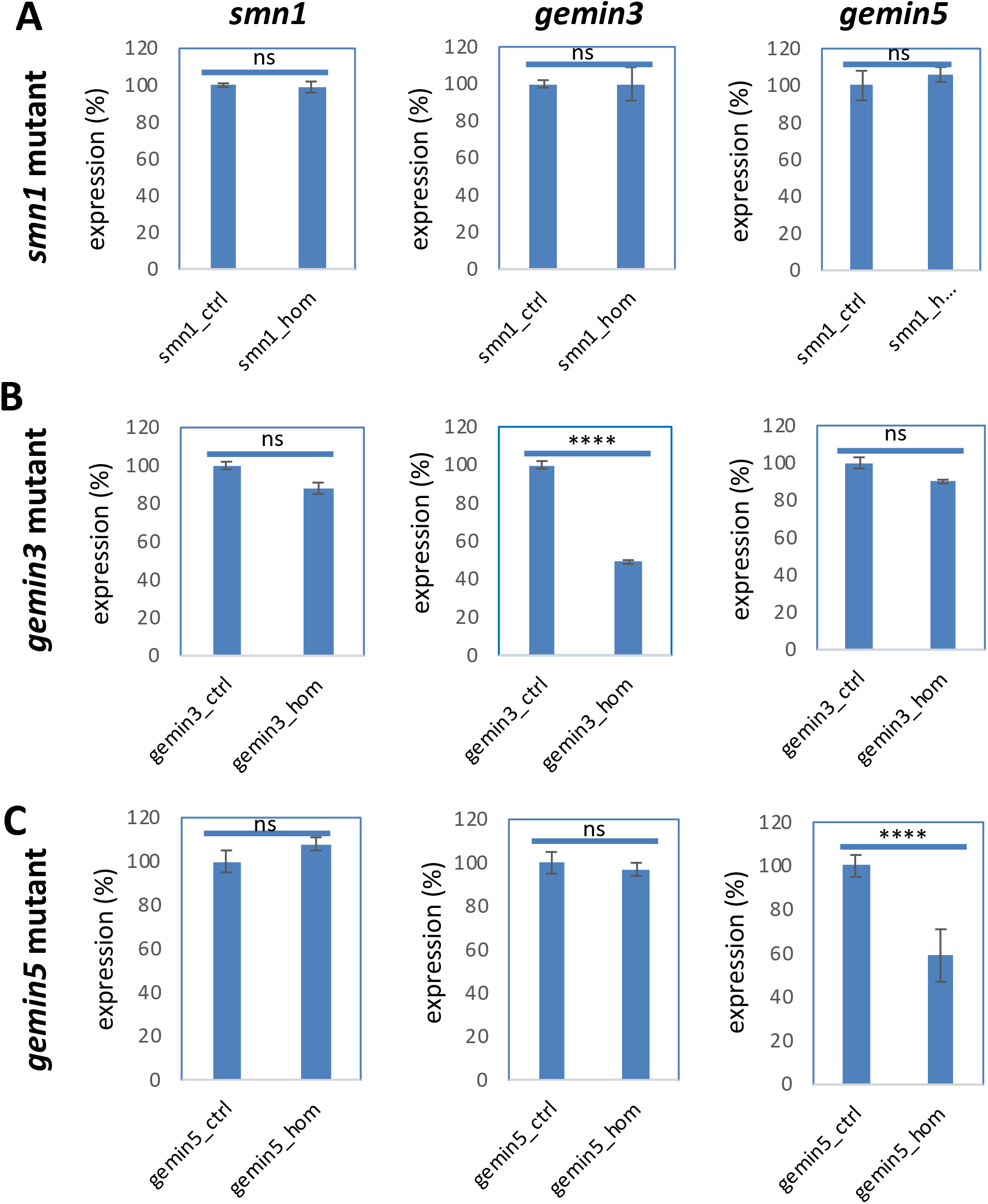
Expression of *smn1, gemin3* and *gemin5* in the control and mutant embryos of *smn1^fh229^* (A), *gemin3^hg105^* (B) and *gemin5^hg80^* (C). Genes examined are labeled on the top. Mutants used are labeled on the left. Graphs show the mean ± s.e.m. ns, P > 0.05; ****P < 0.0001.

**Suppl. Fig. 9.**
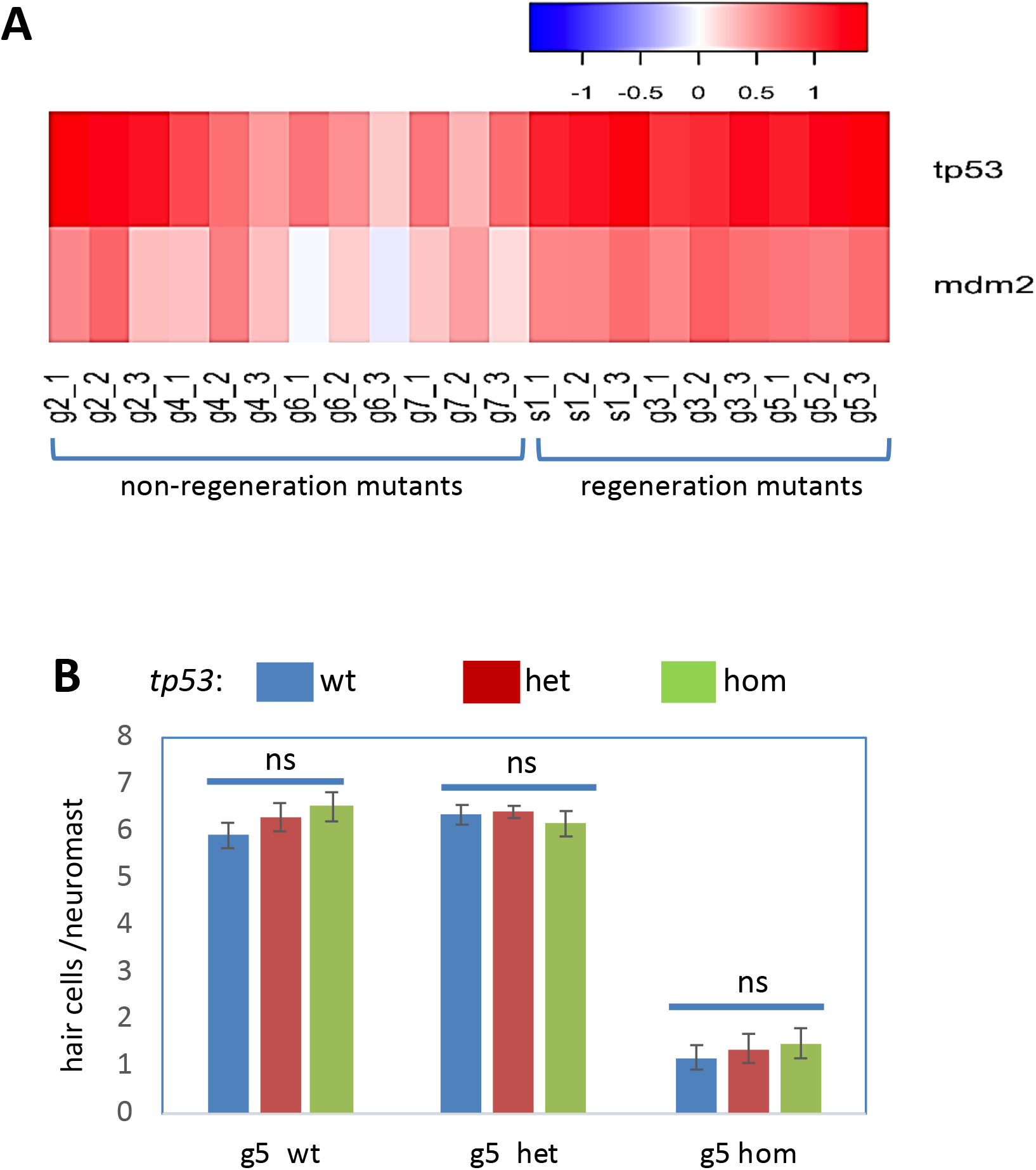
*tp53* knockdown had no impact on regeneration in the *gemin5* mutant. (A) Heat map of *tp53* and *mdm2* expression in the RNA-sequencing samples. (B) Hair cell regeneration analysis for the *tp53^zdf1^*/*gemin5^hg81^* double mutants. Graph shows the mean ± s.e.m. The *tp53^zdf1^* mutation caused no difference in the wild-type (wt), heterozygotes (het), or homozygotes (hom) of *gemin5* embryos (ns, P > 0.05).

**Suppl. Table 1.** Genetic mutations used in this study.

| Genes | Mutations | References |
| --- | --- | --- |
| *smn1* | Y262X | fh229, (54) |
| *smn1* | 2 bp insertion | hg104, this study |
| *gemin3* | 10 bp deletion | hg105, this study |
| *gemin3* | 9 bp deletion | hg106, this study |
| *gemin5* | 2 bp deletion | hg107, this study |
| *gemin5* | 1 bp deletion | hg80, (10) |
| *gemin5* | 20 bp deletion | hg81, (10) |
| *gemin2* | 7 bp deletion | hg108, this study |
| *gemin4* | 11 bp insertion | hg109, this study |
| *gemin6* | 14 bp deletion | hg110, this study |
| *gemin7* | 1 bp deletion | hg111, this study |
| *gemin8* | 7 bp insertion | hg112, this study |
| *strap/unrip* | 8 bp deletion | hg113, this study |
| *nrg1* | 10 bp deletion | hg114, this study |
| *erbb3b* | 7 bp deletion | hg115, this study |
| *tp53* | M214K | zdf1, (30) |
| *tp53* | 7 bp deletion | hg91, (31) |

**Suppl. Table 2.** Mutations in seven of the nine SMN complex members affect adult survival.

| Genes | Mutations | Fish # _total | Fish#_wt | Fish#_het | Fish#_hom |
| --- | --- | --- | --- | --- | --- |
| *smn1* | Y262X | 20 | 7 | 13 | 0 |
| *gemin2* | 7 del | 19 | 7 | 12 | 0 |
| *gemin3* | 9 del | 20 | 8 | 12 | 0 |
| *gemin4* | 11 ins | 27 | 10 | 17 | 0 |
| *gemin5* | 1 del | 25 | 8 | 17 | 0 |
| *gemin6* | 14 del | 34 | 9 | 25 | 0 |
| *gemin7* | 1 del | 26 | 9 | 17 | 0 |

**Suppl. Table 3.**
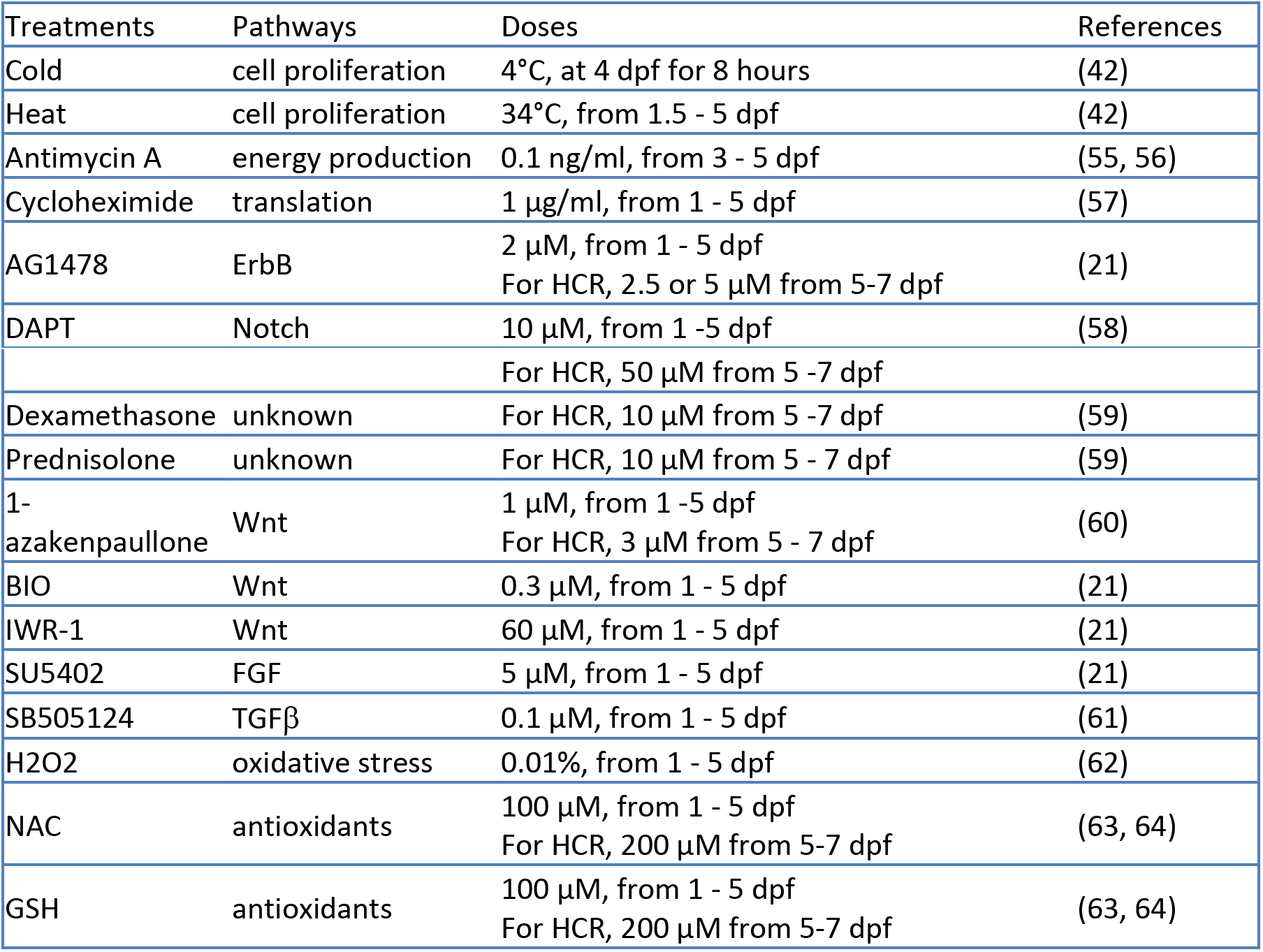
Treatments used for screening the biological pathways that contribute to the regeneration phenotype.

